# Trained Immunity Affecting Dendritic Cell Differentiation and Function in Rheumatoid Arthritis

**DOI:** 10.1101/2025.02.28.640673

**Authors:** Lin Tze Tung, Roselyn R. Jiang, Bianca Pozzebon, Mathieu Mancini, Lucie Droin, Mitra Yousefi, Joo Eun (June) Kim, Marwah Mousa, Yue Liang, Connie Krawczyk, Ines Colmegna, Danielle Malo, David Langlais, Silvia M. Vidal, Anastasia Nijnik

**Affiliations:** Department of Physiology, McGill University, Montreal, QC, Canada; McGill University Research Centre on Complex Traits, McGill University, QC, Canada; Dahdaleh Institute of Genomic Medicine, Montreal, QC, Canada; Department of Microbiology and Immunology, McGill University, Montreal, Canada; Department of Human Genetics, McGill University, Montreal, QC, Canada; Department of Metabolism and Nutritional Programming, Van Andel Institute, Grand Rapids, MI, USA; Division of Rheumatology, Department of Medicine, McGill University Health Centre (MUHC), Montreal, QC, Canada; The Research Institute of the McGill University Health Centre (MUHC), Montreal, QC, Canada; Department of Medicine, McGill University, Montreal, QC, Canada

**Author notes:** **Co-corresponding authors:** Dr. Anastasia Nijnik, Bellini Life Sciences Complex, Rm 368, 3649 Promenade Sir William Osler, H3G 0B1 Montreal, QC, Canada. Dr. Silvia Vidal, Bellini Life Sciences Complex, Rm 367, 3649 Promenade Sir-William-Osler, H3G 0B1, Montreal, QC, Canada. Dr. David Langlais, Dahdaleh Institute of Genomic Medicine, 740 Ave Dr Penfield, Rm 4203, H3A 0G1 Montreal, QC, Canada.

**Keywords:** rheumatoid arthritis, dendritic cells, trained immunity, emergency myelopoiesis, inflammation, mouse models

## Abstract

Rheumatoid arthritis affects ∼0.5-1% of the adult population and results in joint inflammation, chronic pain, and many systemic comorbidities. Immune and inflammatory tissue damage is the main pathogenic mechanism in rheumatoid arthritis, and involves the hyperactivation of both innate and adaptive immune systems. Trained immunity has become well established as an important feature of the innate immune system that allows the host to mount functionally altered immune responses based on their previous history of immune exposures, independently of the classical adaptive immunological memory. However, the role of trained immunity in systemic autoimmune and inflammatory disorders remains poorly understood. In the current work, we demonstrate that emergency myelopoiesis is induced in chronic rheumatoid arthritis in murine models and acts not only to enhance innate immune cell numbers but also to produce functionally altered innate immune cells. Such effects are cell-intrinsic to hematopoietic stem and progenitor cells (HSPCs) and persist independently of the inflammatory disease milieu. Importantly, these trained immunity mechanisms impact not only macrophages but also dendritic cells, which are the major antigen presenting cells that bridge the innate and adaptive immune responses. ‘Trained’ dendritic cells show changes in global gene expression profiles, and significantly altered responses to recall immune stimulation and capacity for T cell activation. This study therefore represents the first demonstration of ‘dendritic cell trained immunity’ in rheumatoid arthritis.

## Introduction

Rheumatoid arthritis is a chronic autoimmune and inflammatory disease that affects ∼0.5-1% of the adult population, with the incidence of the disease elevated among women, smokers, and first-degree relatives, as well as strongly increased with age (1, 2). While joint inflammation is the primary feature of the disease, many rheumatoid arthritis patients develop extra-articular manifestations and associated comorbidities, such as hypertension, osteoporosis, interstitial lung disease, and others (3, 4). Overall, rheumatoid arthritis is chronic and progressive, causing pain, impaired mobility, reduced quality of life, and premature mortality. Despite significant progress in the available therapeutic options, a cure for rheumatoid arthritis is still lacking (5).

Immune and inflammatory tissue damage is the primary mechanism for arthritis disease onset and progression, with a hyperactivation of both innate and adaptive immune systems, release of inflammatory mediators, influx of activated monocytes and other myeloid leukocytes into the affected tissues, and a systemic loss of B and T cell tolerance (6, 7). While these and other immunological disease mechanisms in rheumatoid arthritis are widely investigated, it is essential to consider that most immune cells are short-lived and rapidly replenished from hematopoietic stem and progenitor cells (HSPCs) in the process of hematopoiesis (8). Hematologic dysfunction is common in arthritis patients, with anemia and clonal hematopoiesis being most widely studied (9, 10). We argue that the full understanding of immune dysfunction in rheumatoid arthritis and its roles in the disease pathophysiology requires not only the analyses of peripheral terminally differentiated immune cells, but also the understanding of the mechanisms of immune cell production, differentiation, and turnover.

Over the past decade, trained immunity has become established as an important feature of the innate immune system that allows the host to mount functionally altered immune responses based on their previous history of immune exposures, independently of adaptive immune memory (11, 12). While some long-lived tissue-resident innate immune cells can act as mediators of trained immunity, we and others have shown that altered HSPC function through their long-term epigenetic, transcriptional, and metabolic reprogramming is the major carrier and mediator of trained immunity in murine models of Bacillus Calmette–Guérin (BCG) vaccination, various infectious diseases, and other pathologies (13, 14, 15, 16, 17). There is therefore a growing interest in the role of trained immunity in systemic autoimmune and inflammatory disorders, including rheumatoid arthritis, although this remains poorly understood (18, 19).

Systemic hematologic dysfunction, with an induction of emergency myelopoiesis, is reported in recent studies in murine models of rheumatoid arthritis and related disorders, and has been shown to contribute to the disease pathogenesis (20, 21). Furthermore, trained immunity established in HSPCs in murine periodontitis increases the severity of collagen antibody-induced arthritis in mice (CAIA) (22), indicating that at least in some settings trained immunity is capable of modulating arthritis disease progression. In human rheumatoid arthritis patients, we characterized cell-intrinsic changes in HSPC functions, including altered cell numbers, impaired ex vivo proliferation, altered signaling pathway activities, and accelerated telomere loss (23, 24, 25). Furthermore, monocytes from arthritis patients mount enhanced inflammatory responses following ex vivo restimulation (18, 19), however it is unclear if such changes in monocyte functions are long-term and constitute a true form of immunological memory. In summary, rheumatoid arthritis is associated with altered HSPC function and an induction of emergency myelopoiesis to ramp up innate immune cell production. However, the cell-intrinsic impact of these effects not only on the innate immune cell numbers but on the functional properties of the innate immune cells, independently of the inflammatory disease milieu, remains poorly understood.

Dendritic cells are the major professional antigen presenting cells of our immune system, defined by their capacity to activate naive T cells and therefore induce antigen-specific adaptive immune responses (26). Based on their ontogeny in the bone marrow and functional properties, dendritic cells are broadly classified as conventional (cDCs) and plasmacytoid (pDCs), with monocyte-derived dendritic cells (moDCs) developing in the periphery and sharing some properties of cDCs (26). Importantly, beyond their immunogenic functions, dendritic cells play essential but highly complex roles in regulating T cell immune response polarization (27), and as mediators of immune tolerance through the induction of anergic and regulatory T cells (Tregs) (28, 29). Dysregulation of dendritic cell function is therefore an important component in the pathogenesis of autoimmune disorders, including rheumatoid arthritis. Notably, dendritic cells are short-lived, with an estimated lifespan of 10-30 days in mice (30, 31). They are therefore rapidly replenished from HSPCs, with cDCs arising from the common myeloid progenitor (CMPs) together with monocytes and other myeloid leukocytes (32). Indeed, induction of dendritic cell development is reported in infectious diseases (33, 34, 35) and investigated as a form of immunotherapy for cancer (36). Furthermore, vaccination against the opportunistic fungal pathogen *Cryptococcus neoformans* (37) or an intradermal inoculation with cholera toxin B subunit (38) in murine models result in long-term changes in dendritic cell function. Overall, this indicates that trained immunity can impact dendritic cell function in some disease settings. However, in chronic inflammatory conditions such as rheumatoid arthritis the cell intrinsic impact of the disease on dendritic cells, including their capacity to respond to microbial stimulation and to activate T cell responses, remains unknown.

In the current study, we demonstrate that emergency myelopoiesis induced in chronic rheumatoid arthritis acts not only to enhance innate immune cell numbers but results in the production of functionally altered innate immune cells. Such effects are cell-intrinsic to hematopoietic stem and progenitor cells (HSPCs) and persist independently of the inflammatory disease milieu. Importantly, these trained immunity mechanisms impact not only macrophages but also dendritic cells, altering their global transcriptional profiles, responses to immune stimulation, and capacity for T cell activation. This study therefore represents the first demonstration of ‘dendritic cell trained immunity’ in rheumatoid arthritis.

## Materials and Methods

### Mouse Experiments

SKG mouse line carries a homozygous G-to-T substitution at nucleotide 489 in the *Zap70* gene on the BALB/c genetic background (39). Rheumatoid arthritis disease was induced in the mice with an intraperitoneal injection of zymosan (Millipore-Sigma, 58856-83-2) at 5 mg per mouse at the age of 8-10 weeks, as previously described (40). Control mice received an equal volume of PBS as vehicle control. The mice were analyzed at ≥5 weeks after the zymosan treatment when the animals had a clinical score of 8-16, indicating medium to severe arthritis (41). Collagen-induced arthritis (CIA) mouse model was established on genetically susceptible DBA/1 genetic background with an intradermal immunization of chicken type-II collagen (100 μg, Chondrex, 20011) emulsified in an equal volume of complete Freund’s adjuvant containing 4 mg/mL of heat-denatured *Mycobacterium* (Chondrex, 7001) administered at the base of the tail (40, 41). Mice were maintained under specific pathogen free conditions, and matched by age and sex between the experimental groups. All work was in accordance with the guidelines of the Canadian Council on Animal Care and protocols MCGL 7932 and 6029 approved by the McGill Animal Care Committee.

### ARRIVE Animal Experimentation Statement

Mouse experiments are reported according to the ARRIVE guidelines. Experimental unit corresponds to a single animal. Studies used both female and male mice, sex-matched between experimental groups, however most animals were female to reflect the stronger predisposition of this sex to arthritis in patients and mouse models. Test and control mice were bred as littermates and maintained in shared cages. The mice were 8-10 weeks of age at the time of disease induction. Staff were not blinded to group allocations, and no *a priori* sample size calculations were carried out.

### Flow Cytometry

Cell suspensions of mouse tissues were prepared in RPMI-1640 (ThermoFisher Scientific) with 2% (v/v) fetal calf serum (FCS, ThermoFisher Scientific), 100μg/ml streptomycin (Wisent), and 100U/ml penicillin (Wisent). The samples were stained for cell surface markers in PBS with 2% (v/v) FCS and Super Bright Complete Staining Buffer (SB-4401-75, 1:20, ThermoFisher Scientific) for 20 minutes on ice, using the fluorophore conjugated antibodies provided in Supplemental Table S1. Viability Dye eFluor 506 (ThermoFisher Scientific) was used to discriminate dead cells. Compensation was performed with BD CompBeads (BD Biosciences) or UltraComp eBeads Plus (ThermoFisher Scientific). The data were acquired on BD Fortessa and analyzed with FlowJo software (TreeStar, BD Biosciences).

### Bone Marrow Derived Macrophage Cultures

Murine bone marrow cells were resuspended at 10^7^ per mL and seeded dropwise at 2 mLs per plate into the 150 x 25 mm non-tissue culture treated plates (Fisher Scientific) with 25 mL of complete DMEM media, containing 10% FCS (both from ThermoFisher Scientific), 30% L-cell conditioned media (produced from L929 cell line) and 100 U/mL penicillin/streptomycin (Wisent). All cultures were fed with fresh media at day 4 and harvested on day 7.

### Bone Marrow Derived Macrophage Stimulations

Murine bone marrow derived macrophages (BMDMs) were harvested on day 7 and seeded at 2×10^5^ cells per well into non-tissue culture treated 96 well plates (Falcon) with 200 μl of complete DMEM media to rest in an incubator at 37°C and 5% CO_2_ overnight. Stimulations were performed next day with lipopolysaccharide (LPS, *E. coli* O111:B4, Millipore-Sigma, 10 ng/mL), polyinosinic–polycytidylic acid (poly(I:C), Millipore-Sigma, 20 μg/mL), (1→3)-β-D-glucan from *Alcaligenes faecalis* (Millipore-Sigma, 5 μg/mL) or blank media as negative control for 12 hours. Supernatants were collected for cytokine analyses, and BMDMs prepared for flow cytometry, staining with Fc blocker (TruStain FcX anti-mouse CD16/32 antibody, BioLegend #101320) and viability dye (Fixable Viability Dye eFlour 506, Thermo Fisher Scientific, #65-0866-14). For analyses of cell surface markers, BMDMs were stained with the fluorophore conjugated antibodies listed in Supplemental Table S2. For analyses of cytokine production, BMDMs were pre-treated with Monensin (BioLegend #420701) and Brefeldin A (BioLegend #420601) at the manufacturer’s recommended concentration (1:1000) for 4 hours, and processed with BD Fixation/ Permeabilization Solution Kit (BD Biosciences #554714) prior to intracellular staining. For the analyses of phagocytic activity, pHrodo Red *E. coli* BioParticles Conjugates (ThermoFisher Scientific, P35361) were used according to the manufacturer’s protocols.

### Bone Marrow Derived Dendritic Cell Cultures

Murine bone marrow derived dendritic cells (BMDCs) were derived as previously described (42, 43). Bone marrow cells were resuspended at 10^7^ per mL and seeded dropwise at 75 μLs per well into 6-well non-tissue culture treated plates (Fisher Scientific) containing 3 mL/well of RPMI-1640 media, with 10% FCS (both from ThermoFisher Scientific), 2 mM L-glutamine (Wisent), 100 U/mL penicillin/streptomycin (Wisent), 50 μM β-mercaptoethanol (Millipore-Sigma), and 20 ng/mL granulocyte-macrophage colony-stimulating factor (GM-CSF, #315-03, PeproTech, ThermoFisher Scientific) (44). In many experiments, magnetic depletion of mature immune cells from the bone marrow samples was performed using the EasySep Streptavidin RapidSpheres Isolation Kit (#19860, Stem Cell Technologies) and biotin-conjugated antibodies against the major blood and immune lineage markers: CD3 (BioLegend #100244, 17A2), CD19 (BioLegend #115504, 6D5), NK1.1 (BioLegend #108704, PK-136), TER119 (BioLegend #116204), Ly6G (BioLegend #127604, 1A8), and F4/80 (BioLegend #123106). The resulting lineage depleted murine bone marrow cells were resuspended at 10^7^ per mL and seeded dropwise at 70 μLs per well into the same plates and media as described above. All BMDC cultures were fed with fresh media at days 3 and 6.

### Bone Marrow Derived Dendritic Cell Stimulation

BMDCs were harvested at days 8-9, counted, and seeded at 10^6^ cells/mL for stimulation with lipopolysaccharide (LPS, *E. coli* O111:B4, Millipore-Sigma, 10 ng/mL), polyinosinic–polycytidylic acid (poly(I:C), Millipore-Sigma, 20 μg/mL), or (1→3)-β-D-glucan from *Alcaligenes faecalis* (Millipore-Sigma, 5 μg/mL) for 16 hours. For the analyses of activation and checkpoint marker expression the cells were stained with fluorophore conjugated antibodies listed in Supplemental Table S3. For intracellular flow cytometry analyses of cytokine production, Brefeldin A and Monensin were added to block cytokine secretion over the last 4 hours of stimulation (BioLegend #420701, #420601, 1:1000), BMDCs were fixed and permeabilized with the BD Fixation/Permeabilization Kit (BD Biosciences, #554714), and stained with PE anti-mouse IL-12/IL-23 p40 antibody (BioLegend #505204).

### Antigen Presentation and T cell Stimulation Assays

The assays were performed as previously described (42). Briefly, BMDCs were either untreated or stimulated with LPS (10 ng/ml, *E. coli* O111:B4, Millipore-Sigma), poly(I:C) (Millipore-Sigma, 20 μg/mL), or β-D-glucan (Millipore-Sigma, 5 μg/mL), together with whole ovalbumin (OVA, endotoxin-free, 0.25 mg/mL, Worthington, LS003061) for 16 hours. OT-II CD4 T cells were isolated from the spleens of C.Cg-Tg(DO11.10)10Dlo/J mice (Jackson Laboratories, JAX 003303) using the EasySep Mouse CD4^+^ T Cell Isolation Kit (Stem Cell Technologies, #19852). In some assays, T cells were pre-loaded with 10 µM Cell Proliferation Dye eFluor 450 (ThermoFisher Scientific), according to the manufacturer’s protocol. BMDCs were co-cultured with OT-II CD4 T cells in round-bottom 96-well plates for 3-4 days, in IMDM media with 10% FCS (ThermoFisher Scientific), 2 mM L-glutamine (Wisent), 100 U/mL penicillin/streptomycin (Wisent), and 50 μM β-mercaptoethanol (Millipore-Sigma), at 10^4^ BMDCs and 10^5^ T cells per well (1:10 ratio).

The cultures were analyzed by flow cytometry at day 3, for T cell activation markers CD44 and CD69, and for Cell Proliferation Dye eFluor 450. Alternatively, for the analyses of cytokine production, the cultures were stimulated on day 4 with Cell Activation Cocktail (BioLegend, #423302, 1:500), containing ionomycin and phorbol 12-myristate-13-acetate (PMA) for 4 hours. Brefeldin A and Monensin were added to block cytokine secretion (BioLegend #420701, #420601, 1:1000) over the last 2 hours. The cells were treated with the BD Fixation/Permeabilization Solution Kit (BD Biosciences, #554714), and stained for intracellular cytokines IFNγ, IL-4, IL-17, and IL-10. Alternatively, the cells were treated with the FoxP3 Transcription Factor Staining Buffer Set (ThermoFisher Scientific, #00-5523-00), and stained for transcription factors T-bet, GATA3, RORγt, and FOXP3 and for proliferation marker Ki-67. All flow cytometry analyses were conducted after pre-gating on live CD3^+^CD4^+^ T cells, and the antibodies used are listed in Supplemental Table S4.

### Mouse Bone Marrow Transplantation (or Chimera) Assays

For competitive bone marrow transplantations, cohorts of wild type CByJ.SJL(B6)-Ptprca/J (JAX 006584, congenic for CD45.1) mice were irradiated with 2 doses of 4.5Gy, delivered 3 hours apart, in a RS-2000 irradiator (Rad Source), and engrafted with a 1:1 mix of bone marrow cells from CD45.2^+^ SKG mice with active arthritis (RA-SKG) and from control CD45.1^+^ allotype-marked mice, delivered as intravenous injection in sterile PBS. The mice were kept on neomycin in drinking water (2g/l, BioShop) for 3 weeks. Some cohorts of the chimeric mice were challenged with an intraperitoneal injection of zymosan (Millipore-Sigma, 58856-83-2) at 5 mg per mouse, as described previously (40), prior to the analyses. The analyses were conducted at >20 weeks post-reconstitution, and encompassed comparing the numbers and activation states of CD45.2^+^ versus CD45.1^+^ DCs in lymphoid tissues and in BMDC cultures from the chimeric mice.

### Fluorescence Activated Cell Sorting and RNA Isolation

For the fluorescence activated cell sorting (FACS), BMDCs pre-stimulated in culture for 12 hours were harvested and stained with fluorophore-conjugated antibodies anti-CD11c APC (Biolegend, N418, #117310), anti-CD11b PerCP-Cy5.5 (Thermo Fisher, M1/70, #45-0112-82), and with the viability dye DAPI (BioLegend). Cell sorts were performed on the FACSAria Fusion (BD Biosciences) into the Mag-MAX Lysis Buffer (Ambion, AM1830). RNA was isolated using the Mag-MAX total RNA kit (Ambion, AM1830) according to the manufacturer’s protocol, and RNA yields and quality were assessed on Bioanalyzer (Agilent).

### Myeloid Colony Forming Units Assays

Colony forming units (CFU) assays were performed with primary mouse bone marrow using MethoCult M3534 media following the manufacturer’s protocols (Stem Cell Technologies). Myeloid colonies were counted at 10-12 days of incubation, in duplicates.

### Cytokine Assays

Cytokine levels were analyzed by Eve Technologies Corporation (Calgary, AB, Canada) using the Mouse Cytokine Proinflammatory Focused 10-Plex Discovery Assay Array (MDF10) or the Human Cytokine Proinflammatory Focused 15-Plex Discovery Assay Array (HDF15).

### Hematology analyses

Hematology analyses of mouse blood were performed by the Diagnostics Laboratory of the McGill Comparative Medicine Animal Resources Centre (CMARC), as previously described (45).

### Study subjects

This study was approved by the McGill University Health Center Ethics Review Board (Protocol number MP-37-2021-7495). Patients who fulfilled the following criteria were included: age >18, rheumatoid arthritis classification according to the 2010 American College of Rheumatology/European League Against Rheumatism criteria, seropositive for rheumatoid factor and/or ACPA, and no coexisting systemic inflammatory disorders, infectious diseases, or cancer. Healthy controls age- and sex-matched to rheumatoid arthritis patients were included. All participants provided written informed consent. The demographic characteristics of study participants are summarized in Supplementary Table S5.

### Human monocyte-derived dendritic cells

Peripheral blood mononuclear cells (PBMCs) were isolated from 40 mL of peripheral blood by gradient centrifugation using Lymphoprep and SepMate-50 IVD tubes (Stem Cell Technologies, #7811, #85450). Subsequently, monocytes were isolated with the EasySep Human Monocyte Enrichment Kit (Stem Cell Technologies, #19058), according to the manufacturer’s protocols. Monocytes were resuspended at 1×10^6^ cells per mL in RPMI-1640 (ThermoFisher Scientific) with 10% (v/v) fetal calf serum (ThermoFisher Scientific), 100μg/ml streptomycin (Wisent), 100U/ml penicillin (Wisent), GM-CSF (100 ng/mL, ThermoFisher, PeproTech 300-03), and IL-4 (50 ng/mL, ThermoFisher, PeproTech 200-04). Monocytes were plated at 3 mLs per well into non-tissue culture treated 6-well plates, as a homogeneous cell suspension. The cell cultures were fed with fresh media on days 3 and 6, and harvested for stimulation on day 7. Stimulation was performed as with murine DCs, using lipopolysaccharide (LPS, E. coli O111:B4, Millipore-Sigma, 10 ng/mL), polyinosinic–polycytidylic acid (poly(I:C), Millipore-Sigma, 20 μg/mL), or (1→3)-β-D-glucan from Alcaligenes faecalis (Millipore-Sigma, 5 μg/mL) for 16 hours. The cells were analyzed by flow cytometry using the antibodies listed in Supplemental Table S6.

### RNA Sequencing

Bulk RNA-seq was conducted as previously described (46, 47). rRNA depletion and library preparation were performed using the KAPA RNA HyperPrep Kit with RiboErase (Roche). Libraries were sequenced on Illumina Novaseq 6000 in a paired-end 100 bp configuration aiming for 50×10^6^ reads per sample. The quality of sequencing reads was validated using FastQC tool (Babraham Bioinformatics), and low-quality bases were trimmed from read extremities using Trimmomatic v.0.36 (48). Sequencing reads were mapped to the mouse UCSC GRCm38/mm10 reference genome assembly using Hisat2 v2.2.1 (49, 50, 51). Gene expression was quantified by counting the number of uniquely mapped reads with featureCounts using default parameters (Subread package v1.5.2) (52). Residual rRNA reads were removed, and only genes expressed above 5 counts per million reads (CPM) in at least 2 samples were retained, for a total of 11,307 genes. Dimension reduction analysis was performed using the Principal Component and Partial Least Squares regression method (53). TMM normalization and differential gene expression analyses were conducted using the edgeR Bioconductor package (54). Genes with changes in expression ≥ |2.0| fold and Benjamini-Hochberg adjusted *p* values ≤ 0.001 were considered significant. Gene Set Enrichment Analyses (GSEA) were performed with GSEA v4.3.2 using MSigDB v2022.1 with default configuration and permutation within the gene sets (55), and Gene ontology (GO) enrichment analyses on differentially expressed genes were performed with DAVID Bioinformatics Resources 6.8 using the Biological Processes (BP4) collection (56).

### Statistics

Statistical analyses were performed with Prism 7.01-9.5.1 (GraphPad Inc.), using Student’s two-tailed *t*-test to compare two datasets and ANOVA for multiple comparisons; p<0.05 was considered significant.

## Results

### Expansion of HSPCs and induction of emergency myelopoiesis in murine rheumatoid arthritis

SKG mice are a well-established model of rheumatoid arthritis, with the disease characterized as late onset, sustained, and progressive (RA-SKG), thus well reproducing the core features of human arthritis. Mechanistically, SKG mice carry a homozygous mutation in *Zap70* that impairs T cell negative selection (39), however the disease develops only following an intraperitoneal injection of zymosan, highlighting the interplay of innate and adaptive immunity in the disease induction. The disease is characterized by joint swelling in all limbs, loss of body weight, and systemic inflammatory pathologies (57, 58). We have recently validated the induction of arthritis by weeks 4-6 post-zymosan treatment in SKG mice (42).

Hematologic dysfunction and the induction of emergency myelopoiesis in RA-SKG relative to control SKG mice were confirmed in peripheral blood, with a major increase in monocyte and platelet counts, and a reduction in lymphocyte counts (Figure S1A). While there was no significant change in erythrocyte counts, we observed reduced hematocrit, hemoglobin concentration, mean corpuscular volume (MCV), and mean corpuscular hemoglobin concentration (MCHC) in RA-SKG relative to control SKG mice (Figure S1A). Flow cytometry analyses of myeloid cells demonstrated an increase in monocytes in the bone marrow and in monocyte-derived macrophages in both bone marrow and spleen of RA-SKG relative to control SKG mice (Figures S1B-C). Furthermore, the monocytes in RA-SKG mice had an altered activation state, with enhanced levels of activation markers CD40, CD86, MHCII and checkpoint marker PD-L1 (Figures S1D-E). Gating strategies are provided in Figure S2A.

There was a significant expansion of hematopoietic stem and progenitor cells (HSPCs) in the bone marrow of RA-SKG relative to control SKG mice, particularly affecting the latter multipotent progenitors MPP3-4, common myeloid progenitors (CMPs), granulocyte monocyte progenitors (GMPs), and megakaryocyte progenitors (MkPs) (Figure 1A-E). Induction of emergency myelopoiesis was further supported by the increase in myeloid colony forming units (CFU) in the bone marrow of RA-SKG mice (Figure S3A)

**Figure 1.**
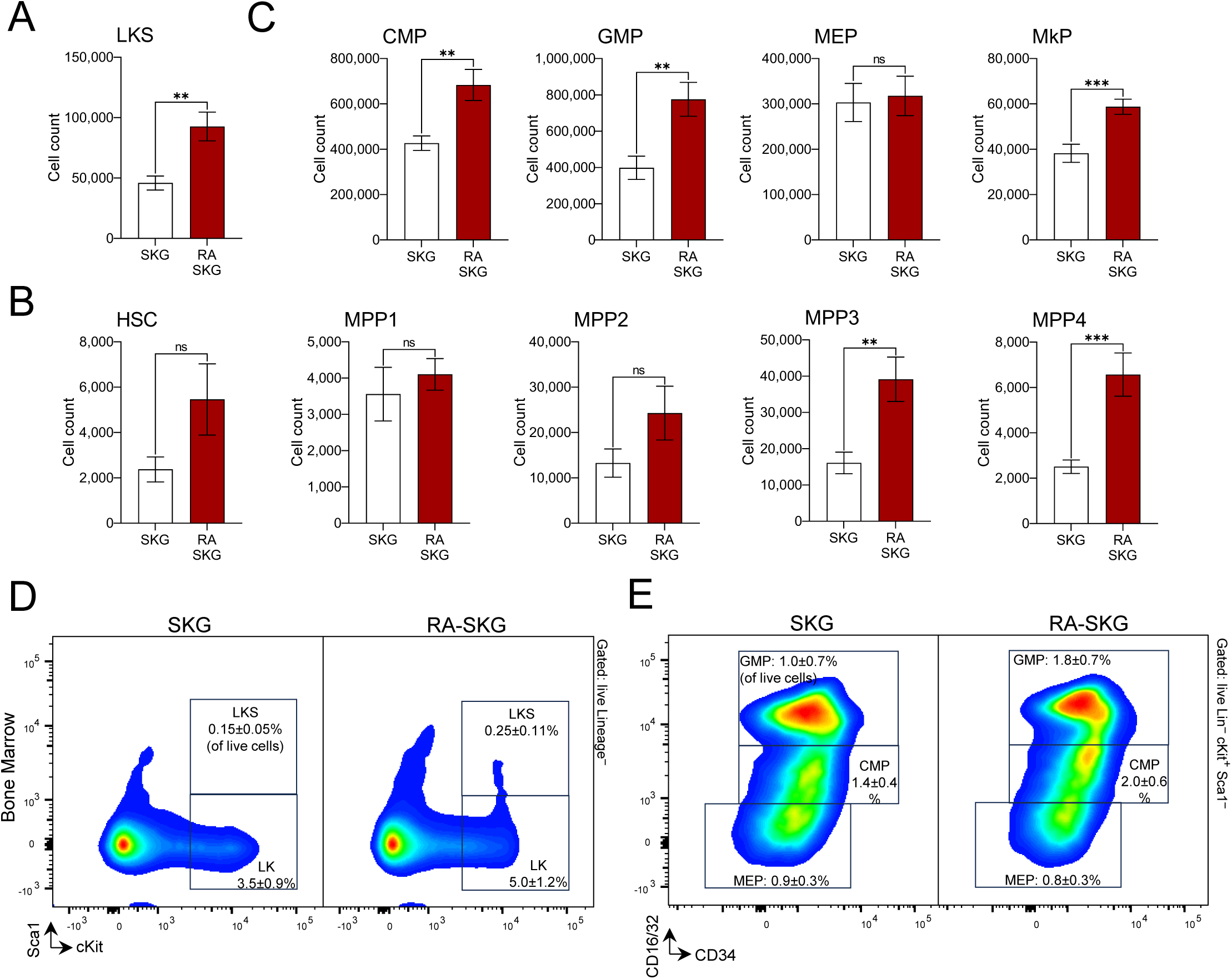
Dysregulation of hematopoiesis in the SKG mouse model of rheumatoid arthritis. **(A)** Increased numbers of LKS hematopoietic stem and progenitor cells in arthritis afflicted SKG mice, gating on Lin^−^cKit^+^Sca1^+^ cells. **(B)** Increased numbers of multipotent progenitor cells MPP3 and MPP4, in arthritis afflicted SKG mice. Cells were gated as Lin^−^cKit^+^Sca1^+^ followed by CD150^+^CD48^−^CD34^−^Flt3^−^ for HSCs, CD150^+^CD48^−^CD34^+^Flt3^−^ for MPP1, CD150^+^CD48^+^CD34^+^Flt3^−^ for MPP2, CD150^−^CD48^+^CD34^+^Flt3^−^ for MPP3, and CD150^−^CD48^+^CD34^+^Flt3^+^ for MPP4. **(C)** Increased numbers of myeloid progenitor cells in arthritis afflicted SKG mice. Cells were gated as Lin^−^cKit^+^Sca1^−^ followed by CD34^+^CD16/32^−^ for common myeloid progenitors (CMP), CD34^+^CD16/32^+^ for granulocyte monocyte progenitors (GMP), CD34^−^CD16/32^−^ for megakaryocyte erythroid progenitor (MEP), and CD41^+^CD150^+^ for megakaryocyte progenitors (MkP). Bars represent means ± SEM; data is from 12-18 mice per group consolidated from 3-4 independent experiments; statistical analyses with unpaired *t*-test; * *p*<0.05, ** *p*<0.01, *** *p*<0.001, ns – not significant; cell counts are presented per two tibias and femurs. **(D)** Representative flow cytometry plots showing the LKS (Lin^−^cKit^+^Sca1^+^) and LK (Lin^−^cKit^+^Sca1^−^), corresponding to HSC-MPPs and early lineage committed progenitors, respectively. **(E)** Representative flow cytometry plots showing the GMP, CMP, and MEP cell populations. Percentage of cells in each gate out of the overall live cell population within the samples is shown as mean ± S.D., consolidating data for 12-18 mice in each group.

Early lineage committed progenitors in RA-SKG mice also showed altered cell surface levels of pattern recognition receptors (PRRs), with the general trend for increased TLR2, CD14 and TREM1 and reduced Dectin-1/CLEC7A (Figure S3B-C). While TLR functions on HSPCs have been widely studied in other settings (59, 60), TREM1 is a poorly characterized PRR induced on myeloid leukocytes under systemic inflammation (61). TREM1 upregulation on myeloid progenitors in arthritis in the current study questions its contribution to the induction of emergency myelopoiesis and whether its functions in HSPCs contribute to its widely established pathogenic roles in arthritis and other inflammatory disorders (62, 63).

In summary, we demonstrate HSPC expansion and induction of emergency myelopoiesis in the SKG murine model of rheumatoid arthritis, mirroring recent studies in related models, including collagen induced arthritis (CIA) (20), collagen antibody induced arthritis (CAIA) (22), and ankylosing spondylitis (21). The altered expression of certain PRRs on the myeloid progenitors in the arthritis-afflicted mice indicates that these cells not only respond to the inflammatory disease milieu but also acquire altered capacity for the sensing of inflammatory and danger signals.

### Ex vivo macrophage differentiation as a model of trained immunity in murine arthritis

To assess if the changes in HSPCs and hematopoiesis in murine arthritis, as reported above, result in the production of functionally altered innate immune cells independently of the inflammatory disease milieu, we applied the bone marrow derived macrophage model (BMDM). This model is widely used to study the mechanisms of both protective and deleterious trained immunity induced with β-glucan, BCG vaccination, or high-fat diet, to give a few examples (16, 17). Bone marrow harvested from RA-SKG and control SKG mice was differentiated ex vivo to BMDMs. The resulting BMDMs were induced with LPS, β-glucan, and poly(I:C), to mimic bacterial, fungal, and viral stimulation, respectively, and analyzed for co-stimulatory and checkpoint markers, for cytokine secretion, iNOS levels, and phagocytic activity. BMDMs from RA-SKG mice showed significantly altered responses to stimulation, with a reduced induction of co-stimulatory markers CD40 and CD86 and an increased induction of checkpoint marker PD-L1 relative to control SKG BMDMs (Figure 2A-B). Furthermore, RA-SKG BMDMs produced less of the pro-inflammatory cytokines IL-1β and IL-6, as well as TNFα under some of the stimulation conditions (Figure 2C), but significantly more of the immunosuppressive cytokine IL-10 (Figure 2C), as shown by analyses of the cell culture supernatants. The reduced production of IL-6 and TNFα by RA-SKG BMDMs was confirmed through intracellular flow cytometry (Figure 2D, S4A-B), further demonstrating their altered responses to immune stimulation. In contrast, the induction of iNOS was significantly enhanced in RA-SKG relative to control BMDMs (Figure S4C-E), while the phagocytic activities of BMDMs measured through pHrodo Red *E. coli* BioParticle uptake were not significantly different (data not shown). In summary, this demonstrates that the inflammatory milieu of murine arthritis triggers cell intrinsic changes in HSPCs and in myeloid cell differentiation to generate macrophages with an altered capacity to respond to immune stimulation. Overall, such macrophages exhibit reduced production of inflammatory cytokines and co-stimulatory molecules and enhanced production of immunoregulatory cytokines and checkpoint markers, as well as enhanced iNOS induction.

**Figure 2.**
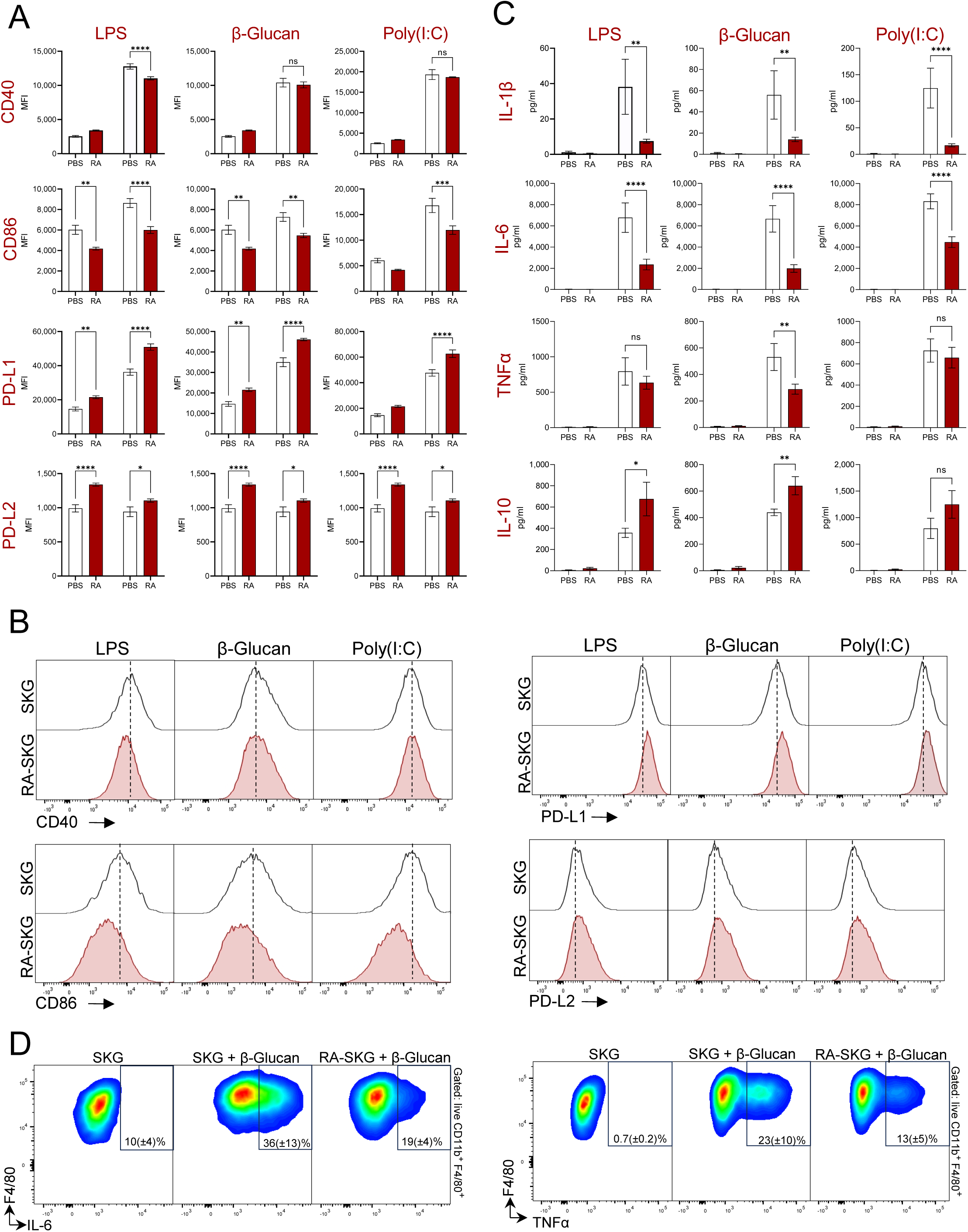
Altered functional profile of macrophages derived *ex vivo* from hematopoietic progenitor cells of mice with active arthritis disease. BMDMs were derived *ex vivo* from the bone marrow of SKG-RA and control SKG-PBS mice, and stimulated with LPS 10 ng/mL, β-glucan 5 μg/mL, or poly(I:C) 20 μg/mL for 12 hours. **(A)** Expression of activation and checkpoint markers on BMDMs, analyzed by flow cytometry, pre-gating on live CD11b^+^F4/80^+^ cells; MFI – mean fluorescence intensity. n=5-6 mice per group, with results reproduced in a second independent experiment. **(B)** Representative flow cytometry histograms showing the expression of co-stimulatory molecules CD86 and CD40, and checkpoint markers PD-L1 and PD-L2 on pre-stimulated BMDMs. **(C)** Analyses of cytokine levels in the BMDM culture supernatants. n=9-11 mice per group, consolidated from two independent experiments. **(D)** Analyses of IL-6 and TNFα production by the BMDMs through intracellular flow cytometry, pre-gating on live CD11b^+^F4/80^+^ cells; n=4 mice per group. Percentages of cells in the positive gates are shown as mean ± S.D. for each group. **(A,C)** Bars represent means ± SEM; statistical analyses with one-way ANOVA and Sidak’s post-hoc test; * *p*<0.05, ** *p*<0.01, *** *p*<0.001.

### Increased dendritic cell numbers and altered activation in murine arthritis

Conventional dendritic cells (cDCs) are short-lived and replenished from common myeloid progenitors (CMPs), like other myeloid leukocytes (30, 31, 32). We therefore tested whether the induction of emergency myelopoiesis in murine arthritis also affects in situ DC numbers. Flow cytometry analyses indicated an expansion of DCs in both bone marrow and spleen of RA-SKG relative to control SKG mice (Figure 3A). Further characterization of DCs as conventional cDCs or plasmacytoid pDCs demonstrated an increase in the numbers of cDC1 cells in the bone marrow and spleen and of cDC2a cells in the bone marrow, while the increases in cDC2a in the spleen and cDC2b in bone marrow and spleen did not reach statistical significance, and pDC numbers were unaffected (Figure 3B). Gating strategies for the analyses are provided in Figure S2B.

**Figure 3.**
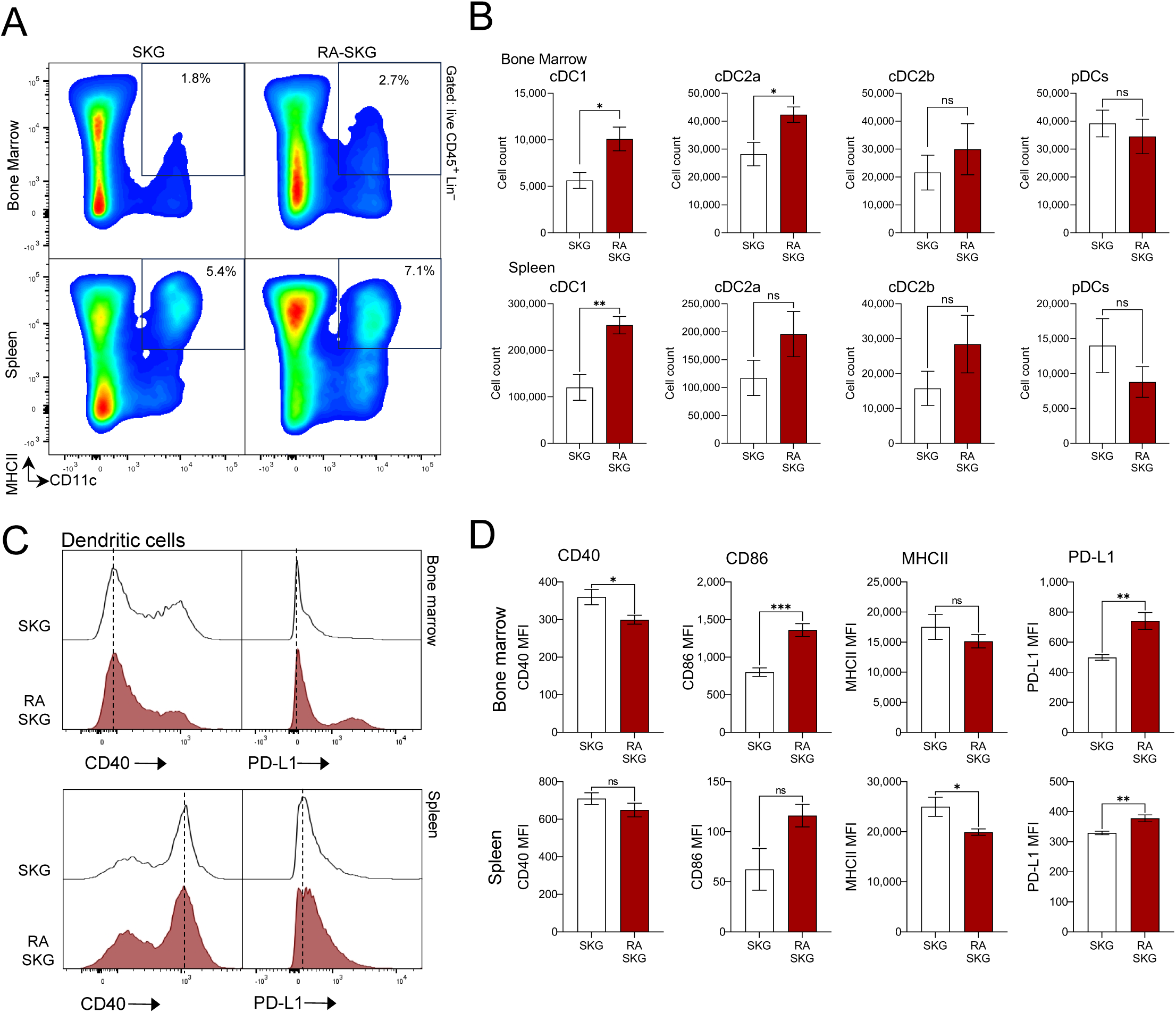
Increased numbers of dendritic cells (DC) and altered DC activation in the SKG mouse model of rheumatoid arthritis. **(A)** Representative flow cytometry plots of bone marrow and spleen from arthritis afflicted RA-SKG and control SKG mice, gated on live CD45^+^Lin^−^ cells and showing the CD11c^+^ MHCII^+^ DC cell population. Full gating hierarchy is presented in Figure S2A; average percentage of cells in DC gates for all mice in each group is indicated. **(B)** Absolute numbers of cDC1, cDC2a, cDC2b, and pDC cells in the bone marrow and spleen of arthritis afflicted RA-SKG and control SKG mice. DCs are defined as live CD45^+^Lin^−^CD11c^+^MHCII^+^ cells, where the lineage markers are CD3, CD19, NK1.1, Ly6G, F4/80, CD64 and TER119. DCs are subdivided into B220^+^PDCA1^+^ pDCs and B220^−^PDCA1^−^ cDCs, which are further subdivided into the cDC1, cDC2a, and cDC2b subsets based on the expression of XCR1, CD172a/SIRPα, and CLEC12A markers, as shown in Figure S2. Bars represent means ± SEM; data are from n=9 mice per group and consolidated from two independent experiments; statistical analyses used Student’s *t*-test; * *p*<0.05, ** *p*<0.01, ns – not significant. Marrow cell counts are per two tibias and femurs. **(C-D)** Expression of activation and checkpoint markers on DCs in the bone marrow and spleen of RA-SKG and control SKG mice, gating on DCs as live CD45^+^Ly6G^−^F4/80^−^CD11c^+^MHCII^+^ cells. Bars represent means ± SEM; data from n=5-9 mice per group; MFI – mean or median fluorescence intensity; statistical analyses with Student’s *t*-test; * *p*<0.05, ** *p*<0.01, *** *p*<0.001; ns – not significant.

The numbers of pre-DCs in the bone marrow of RA-SKG mice also significantly increased (Figure S5A-B), indicating a systemic induction of DC production in murine arthritis. Further flow cytometry analyses of DC surface markers indicated altered DC activation in situ in RA-SKG mice, characterized by increased PD-L1 in the bone marrow and spleen, reduced CD40 in the bone marrow, reduced MHCII in the spleen, and increased CD86 in the bone marrow (Figure 3C-D). Overall, we conclude that the induction of emergency myelopoiesis in murine arthritis is associated with increased DC abundance and altered DC activation.

### Cell intrinsic changes in DC development and function in murine arthritis

cDCs derive from the same early myeloid precursors as monocytes; however, while the induction of trained immunity is widely studied in monocytes and monocyte-derived macrophages in various disease states, its effects on DCs remain largely unexplored. To characterize the cell intrinsic changes in DC development and function in murine arthritis, independently of the in vivo inflammatory disease milieu, DCs were derived ex vivo from the bone marrow of RA-SKG and control SKG mice in GM-CSF supplemented cultures (BMDCs) (44). DC responses to immune stimulation were then analyzed through cytokine production and cell surface marker expression, according to the protocols established above for macrophages. Importantly, to focus on the cell-intrinsic aspects of HSPC to DC differentiation, independently of any changes in bone marrow composition and immune cell function in RA-SKG mice, bone marrow samples were magnetically depleted of cells expressing lymphoid, erythroid, granulocyte, and macrophage lineage markers (CD3^+^, CD19^+^, NK1.1^+^, TER119^+^, Ly6G^+^, F4/80^+^), and the BMDC derivation protocols re-optimized for such lineage depleted marrow samples. Lineage depleted marrow efficiently generated BMDCs, with the marrow of arthritis-afflicted mice producing significantly higher BMDC yields (Figure S5C-F). This indicates that the induction of DC production in murine arthritis is at least in part cell intrinsic and independent of inflammatory disease milieu.

DCs derived ex vivo from either total or lineage-depleted bone marrow of RA-SKG mice showed consistent alterations in the induction of costimulatory and checkpoint markers relative to matched DCs from control SKG mice, including reduced CD40 and increased PD-L1 (Figures 4A-B, S6). Supplemental study used wild type BALB/c mice, corresponding to the genetic background of SKG strain. Upon treatment with zymosan according to the same protocols the wild type mice did not develop arthritis and demonstrated that the changes in DC responses to stimulation in SKG were not a side-effect of zymosan treatment but required arthritis disease induction (Figure S7).

**Figure 4.**
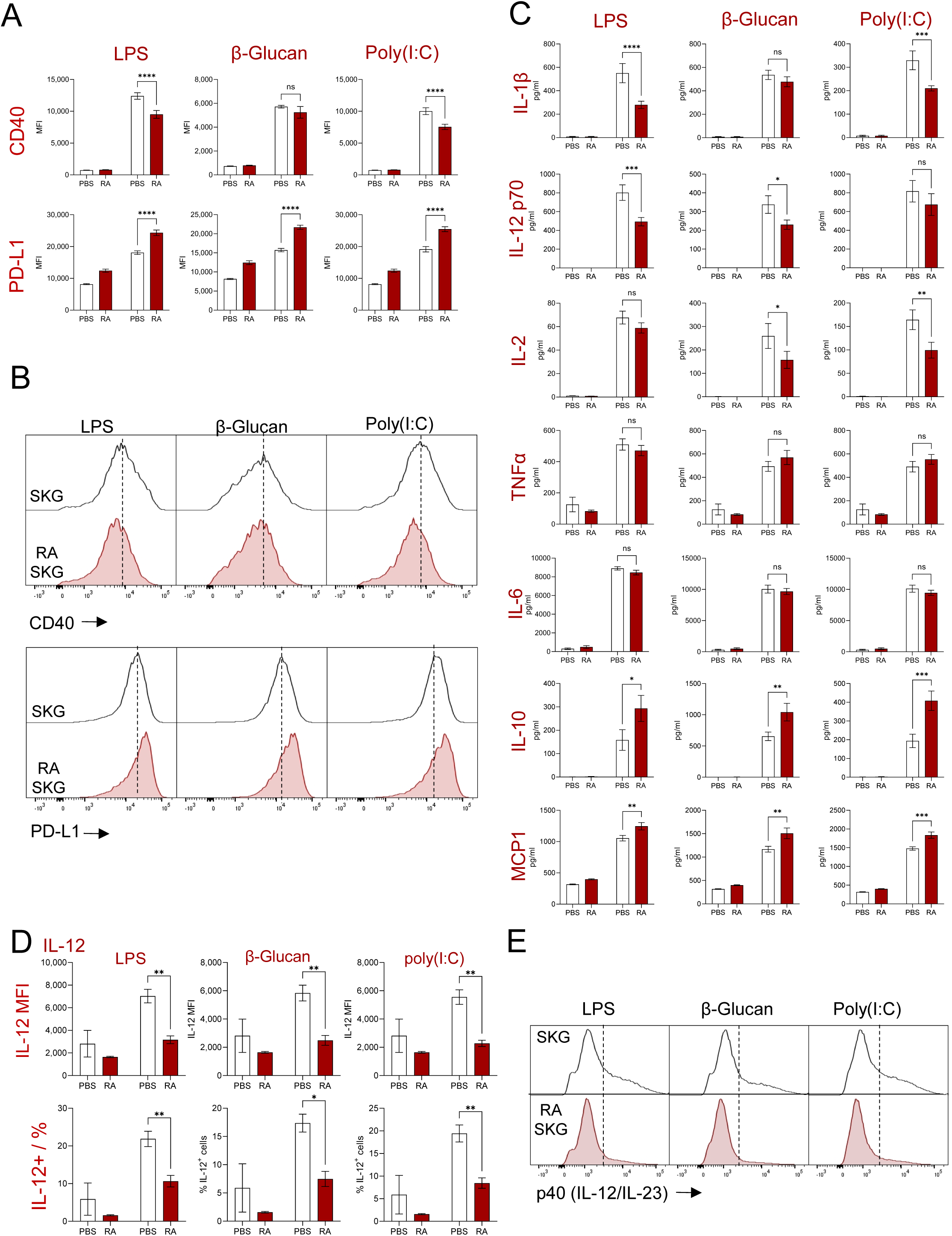
DCs derived *ex vivo* from lineage-depleted marrow of mice with active arthritis disease demonstrate altered activation in response to microbial stimulation. BMDC were derived *ex vivo* from the bone marrow of RA-SKG-RA and control SKG mice, pre-depleted of mature immune cells. BMDCs were stimulated with LPS 10 ng/mL, β-glucan 5 μg/mL, or poly(I:C) 20 μg/mL for 16 hours. **(A)** Expression of activation marker CD40 and checkpoint marker PD-L1 on BMDCs, analyzed by flow cytometry, pre-gating on live CD11b^+^CD11c^+^ cells; MFI – mean fluorescence intensity; n=5-6 mice per group. **(B)** Representative flow cytometry histograms showing the expression of co-stimulatory marker CD40 and checkpoint marker PD-L1 on pre-stimulated BMDCs, gated as live CD11b^+^CD11c^+^ cells. **(C)** Analyses of cytokine levels in the BMDC culture supernatants. Bars represent means ± SEM; statistical analyses with one-way ANOVA and Sidak’s post-hoc test; * *p*<0.05, ** *p*<0.01, *** *p*<0.001; n=5-15 mice per group, and the data are consolidated from 3 independent experiments. **(D-E)** Analyses of BMDC IL-12 production through intracellular staining and flow cytometry, showing **(D)** mean fluorescence intensity (MFI) of IL-12 staining, the percentage of IL-12^+^ cells, and **(E)** representative flow cytometry histograms showing intracellular staining for IL-12, all gated on live BMDCs. Bars represent means ± SEM; statistical analyses with one-way ANOVA and Sidak’s post-hoc test; * *p*<0.05, ** *p*<0.01, *** *p*<0.001; n=4 mice per group.

DCs derived from lineage depleted bone marrow of RA-SKG mice also produced lower levels of inflammatory cytokine IL-1β, lower levels of Th1-polarizing cytokine IL-12, higher levels of immunosuppressive cytokine IL-10, and higher levels of monocyte chemoattractant MCP1/CCL2, with some variation in the magnitude of these effects across the stimulation conditions (Figure 4C). It is interesting to note that reduced IL-1β and increased IL-10 are known to protect against arthritis in vivo in SKG mice (64). In contrast, the production of major inflammatory cytokines TNFα and IL-6 was unchanged between RA-SKG and control DCs (Figure 4C). Intracellular staining and flow cytometry analyses further showed reduced p40 IL-12/IL-23 production in RA-SKG relative to control DCs (Figure 4D-E). As the analyses of DC supernatants showed reduced IL-12 but no detectable IL-23 under any of the conditions (Figure 4C and data not shown), we conclude that this likely reflects a reduction in IL-12 rather than IL-23. Overall, this establishes that the cell intrinsic reprogramming of HSPC to DC differentiation in murine arthritis results in the production of DCs with a significantly altered capacity to respond to immune stimulation.

To test whether the changes in DC development and function described here are specific to the SKG murine model or are a general feature of arthritis, independently of genetic background and disease induction methods, select studies were repeated with the collagen-induced arthritis model (CIA) (41). In this model wild type mice of DBA/1 inbred strain are induced intradermally with type-II collagen in CFA (Col-CFA) and compared to control DBA/1 mice, untreated or treated with CFA alone, as previously described (41) and reproduced in our recent work (42). Col-CFA but not control mouse cohorts developed progressive arthritis from day 18 post-injection and were euthanized for analyses at day 34. DCs derived ex vivo from the bone marrow of arthritis-afflicted Col-CFA mice showed altered expression of costimulatory and checkpoint markers in response to stimulation, with reduced CD40 and enhanced PD-L1 induction (Figure S8), which is fully consistent with our findings from the SKG model. Overall, this confirms the above conclusions and indicates that altered DC development and function in arthritis is conserved across murine models, genetic backgrounds, and disease induction methods.

To test if the altered DC differentiation from the HSPCs of RA afflicted mice can persist long-term, bone marrow cells from CD45.2^+^ SKG mice with active RA (RA-SKG) and control allotype-marked CD45.1^+^ mice were mixed in a 1:1 ratio and transplanted into radioablated CD45.1^+^ recipients. The chimeric mice were analyzed at >20 weeks post-reconstitution, to allow for a full replenishment of the immune system in chimeric mice from the donor HSPCs. Subsequent analyses were conducted either on untreated chimeric mice or on chimeric mice subjected to RA induction via the standard zymosan treatment protocol, analyzing for the relative contribution of CD45.2^+^ RA-SKG versus control CD45.1^+^ HSPCs to the DC lineage. This demonstrated an enhanced contribution of RA-SKG marrow to the DC cell lineage specifically following the zymosan challenge, both in vivo in mouse tissues and ex vivo in the BMDC cultures (Figure S9A-B). Furthermore, RA-SKG DCs from the chimeric mice expressed constitutively higher levels of PD-L1 (Figure S9C). These phenotypes are consistent with the phenotypes of RA-SKG DCs previously observed in the studies above (Figures 3C-D, 4A-B, S6-8), and therefore further support the induction of trained immunity affecting the DC lineage in murine RA.

### Altered DC transcriptional profiles in murine arthritis independent of inflammatory disease milieu

To further understand the cell intrinsic impact of arthritis on DC differentiation and function, independently of the in vivo inflammatory disease milieu, we performed bulk RNA-seq on the BMDCs derived ex vivo from lineage-depleted bone marrow of RA-SKG and control SKG mice. The samples analyzed included BMDCs with and without stimulation, FACS-sorting live CD11c^+^ cells from the cultures for RNA isolation, at n=4-5 mice per group (Figure 5A, Supplemental Table S7).

**Figure 5.**
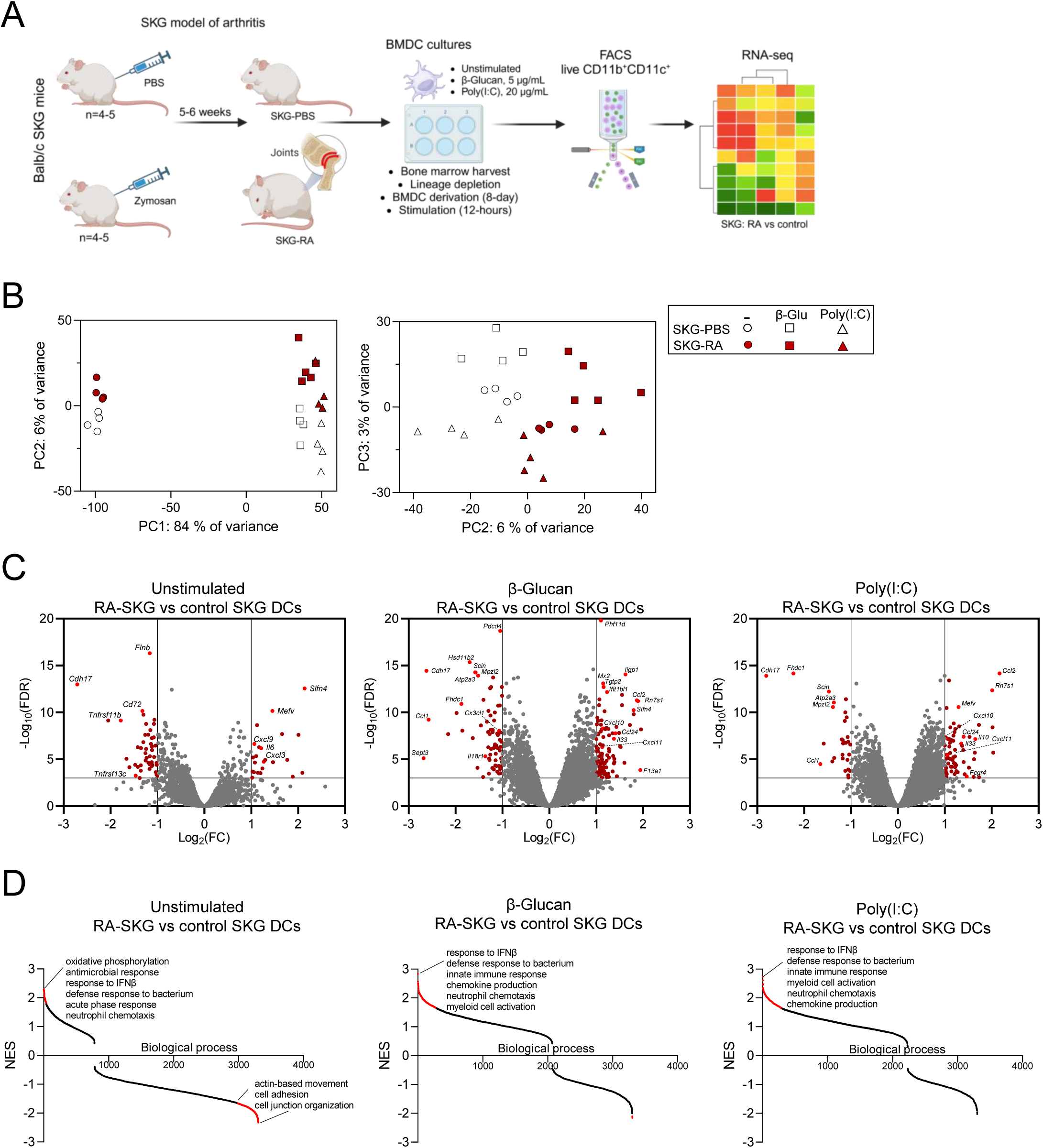
Altered gene expression profiles of DCs derived ex vivo from lineage depleted marrow of arthritis afflicted RA-SKG mice. **(A)** Schematic of the RNA-seq experiment. DCs were derived ex vivo from lineage-depleted marrow of RA-SKG and control SKG mice. The samples analyzed included DCs with and without stimulation, FACS-sorting live CD11b^+^CD11c^+^ cells from the cultures for RNA isolation, at n=4-5 mice per group. **(B)** Dimension reduction analysis of the DC gene expression profiles revealed a clear segregation based on stimulation (PC1 - 84% and PC3 - 3% of variance), and between the RA-SKG and control SKG DCs (PC2 - 6% of variance). **(C)** Volcano plots showing the transcriptional profiles of RA-SKG relative to control SKG DCs, at steady state and following stimulation with β-glucan or poly(I:C). Dysregulated genes (dark red) and dysregulated genes with known immune functions (bright red) are highlighted. **(D)** Gene set enrichment analyses (GSEA) of the transcriptional signatures of RA-SKG relative to control SKG DCs, either unstimulated, or following stimulation with β-glucan or poly(I:C). Normalized enrichment scores (NES) for the pre-established biological process transcriptional signatures are presented. Statistically significant biological processes, based on FDR<0.05, are indicated in red. Select biological processes are labelled on the plot, with the full summary of the GSEA results provided in Supplemental Tables S7E-G.

Dimension reduction analyses of the gene expression profiles revealed a clear segregation of the stimulated and unstimulated DCs (PC1 - 84% of variance), with an induction of classical proinflammatory transcriptional programs, and with some differences between β-Glucan and poly(I:C) stimulation (PC3 - 3% of variance, Figure 5B). Importantly, there was also a clear segregation of the gene expression profiles of DCs derived from arthritis afflicted and control mice, and this segregation was conserved for unstimulated DCs and for DCs following either β-Glucan or poly(I:C) stimulation (PC2 - 6% of variance, Figure 5B). This demonstrates the cell intrinsic impact of the arthritis milieu on bone marrow progenitors to alter the downstream DC development and transcriptional responses to immune stimulation.

Analyzing normalized RNA-seq read counts for the key genes corresponding to the previous readouts in our DC functional assays (Supplemental Table S7A, Figure 4), RA-SKG DCs showed a reduced induction of the gene encoding *Il12b*, and an enhanced induction of *Il10* and *Ccl2* (Figure S10A), fully consistent with our previous cytokine secretion analyses (Figure 4C-E). Furthermore, genes encoding co-stimulatory markers CD80 and CD86 were downregulated and the gene encoding checkpoint marker *Cd274*/PD-L1 was upregulated (Figure S10B), demonstrating the concordance of RNA-seq with previous flow cytometry analyses (Figure 4, S5-7). In contrast, no downregulation of *Il1b* or *Cd40* was observed (Supplemental Table S7A), suggesting that their dysregulation in RA-SKG DCs in previous analyses may be due to post-translational mechanisms.

Differential gene expression analyses comparing DCs derived from the marrow of RA-SKG versus control SKG mice, at fold change (FC) ≥ |2.0|, Benjamini-Hochberg-adjusted *p*-value ≤ 0.001, and read count per million (CPM) > 5 in at least two samples, confirmed the significant shift in the gene expression profile between RA-SKG and control DCs under steady-state unstimulated conditions, with 23 upregulated and 47 downregulated genes (Figure 5C, Supplemental Table S7B). Further transcriptional changes between RA-SKG and control DCs were observed with stimulation, with 109 upregulated and 71 downregulated genes in the β-Glucan condition and 70 upregulated and 30 downregulated genes in the poly(I:C) condition (Figure 5C, Supplemental Tables S7C-D).

Hierarchical clustering of the genes differentially expressed between RA-SKG and control DCs was performed to get a consolidated overview of the transcriptional changes under steady-state and with stimulation. While the transcriptional differences between RA-SKG and control DCs were obvious at steady-state, these differences were mild relative to the transcriptional responses of either group to stimulation (Figure S10C), consistent with previous PCA analyses (Figure 5B). A prevalent pattern emerges whereby the genes upregulated at steady-state in RA-SKG relative to control DCs are either more strongly induced (Cluster 1) or less strongly downregulated (Cluster 3) with stimulation. Accordingly, the genes downregulated in RA-SKG DCs at steady-state are either less strongly induced (Cluster 4) or more strongly downregulated (Cluster 2) following stimulation. Therefore, the overall profile of transcriptional response to microbial stimulation is preserved in RA-SKG DCs, with the same trends in gene upregulation versus downregulation as in the control DCs. However, the magnitudes of the specific stimulation-induced transcriptional signatures in RA-SKG DCs are significantly altered.

Gene set enrichment analyses (GSEA) were conducted comparing RA-SKG and control DCs under steady state unstimulated conditions, as well as following β-Glucan and poly(I:C) stimulation. This demonstrated a constitutive activation of the transcriptional signatures of innate immune and inflammatory response in unstimulated RA-SKG DCs, including ‘antimicrobial response’, ‘response to IFNβ’, ‘type I IFN signaling’, ‘defense response to bacterium’, ‘acute phase response’, and ‘neutrophil chemotaxis’ (Figure 5D, Supplemental Table S7E). Furthermore, the induction of these transcriptional signatures was even more notable when comparing RA-SKG and control DCs following stimulation (Figure 5D, Supplemental Tables S7F-G).

Importantly, the upregulated transcriptional signatures of unstimulated RA-SKG DCs also included ‘oxidative phosphorylation’ and ‘mitochondrial respiratory chain’ (Figure 5D, Supplemental Table S7E), likely indicating that significant metabolic reprogramming may impact RA-SKG DC functions (65). Furthermore, the downregulated transcriptional signatures of unstimulated RA-SKG DCs included ‘cell junction organization’, ‘cell adhesion’, and ‘actin-based movement’ (Figure 5D, Supplemental Table S7E), suggesting possible changes in their migratory capacity. Overall, we conclude that the inflammatory disease milieu makes a stable cell-intrinsic impact on DC differentiation, resulting in altered DC transcriptional profiles at steady state, as well as significantly modulates DC transcriptional responses to microbial stimulation.

### Altered response to microbial stimulation in monocyte derived DCs of rheumatoid arthritis patients

To test if DCs derived *ex vivo* from human rheumatoid arthritis patients also exhibit altered responses to microbial stimulation, monocytes were isolated from peripheral blood of patients and healthy controls matched by sex and age. Monocyte derived DCs (moDCs) were generated over 7 days of culture with GM-CSF and IL-4. Successful derivation of moDCs was confirmed based on the expression of CD11c and DC-SIGN markers (Figure S11A), with no differences in moDC yields between patient and control samples (data not shown). moDCs were stimulated with LPS, β-glucan, or poly(I:C), according to the protocols established above and analyzed for activation markers by flow cytometry. We observed an increased induction of PD-L1 on moDCs from RA patients versus matched healthy controls in responses to β-glucan and poly(I:C), with similar trends observed also for unstimulated and LPS-stimulated moDCs (Figures S11B-C). Although the induction of other activation markers CD40, CD86, and HLA-DR between moDCs of patients and controls was not statistically significant. We also observed reduced secretion of IL-12 p40 in response to β-glucan stimulation, as well as increased secretion of MCP-1 in unstimulated moDCs derived from patients versus healthy controls (Figure S11D-E). Overall, this supports the notion that trained immunity may affect DC differentiation and function not only in mouse RA models but also in rheumatoid arthritis patients, with consistent changes in PD-L1 induction, IL-12 p40, and MCP-1 observed for *ex vivo* differentiated DCs from both RA-SKG mice and human patients.

### Cell intrinsic changes in the capacity of DCs to activate T cells in murine arthritis

The activation of T cells is the primary function of DCs as professional antigen presenting cells. To analyze the impact of arthritis disease on the DC capacity for antigen presentation and T cell activation, independently of inflammatory disease milieu, BMDCs derived ex vivo from RA-SKG and control SKG mice were activated with LPS, β-glucan, or poly(I:C) in the presence of antigenic protein ovalbumin (OVA). DCs were then co-cultured with CD4 T cells expressing a TCR specific for OVA, isolated from OTII transgenic, *Zap70* wild type, BALB/c donor mice. T cells were analyzed at days 3-4 of co-culture, comparing T cell activation by RA-SKG versus control SKG DCs, and measuring cell surface markers of T cell activation, proliferation, and exhaustion, as well as cytokine production, and the expression of transcription factors specific to T-helper and Treg lineages.

CD4 T cell activation in co-cultures with RA-SKG DCs was significantly altered across most of the parameters analyzed. Thus, we observed a significant reduction in IFNγ, IL-4, and IL-17A producing CD4 T cells under some of the stimulation conditions (Figure 6A-B), corresponding to the major Th1, Th2, and Th17 cytokines, respectively. Importantly, there was also a highly significant reduction in IL-10 producing T cells in co-cultures with RA-SKG DCs under all stimulation conditions (Figure 6A-B). Quantification of the average cytokine levels per T cell was also conducted, fully supporting the above conclusions (Figure S12A).

**Figure 6.**
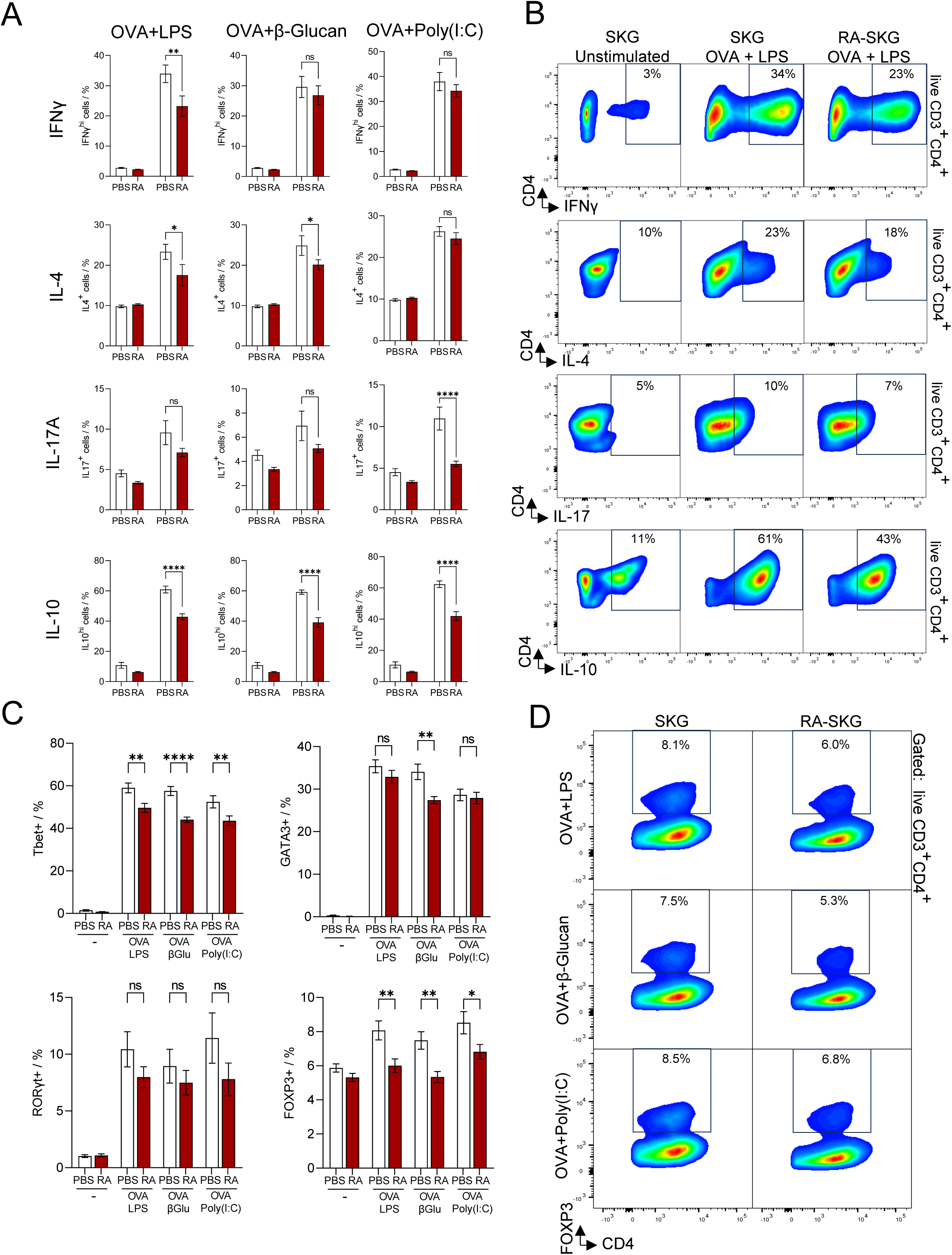
Cell intrinsic impact of arthritis disease milieu on the capacity of DCs for T cell activation. Analyses of OT-II CD4 T cell activation in co-cultures with DCs derived ex vivo from lineage depleted bone marrow of arthritis afflicted SKG-RA or control SKG-PBS mice, pre-stimulated with cognate antigen ovalbumin (OVA), together with LPS 10 ng/mL, β-glucan 5 μg/mL, or poly(I:C) 20 μg/mL for 16 hours. **(A)** Proportion of CD4 T cells in the DC co-cultures positive for cytokines IFNγ, IL-4, IL-17A, and IL-10, analyzed by flow cytometry gating on live CD3^+^CD4^+^ T cells. Positive gates for each channel were set with an FMO control. Bars represent means ± SEM; statistical analyses with one-way ANOVA and Sidak’s post-hoc test; * *p*<0.05, ** *p*<0.01, *** *p*<0.001; n=5-12 mice per group, consolidated from two independent experiments. **(B)** Representative flow cytometry plots showing the IFNγ^+^, IL-4^+^, IL-17^+^, and IL-10^+^ CD4 T cells in the DC co-cultures, analyzed by flow cytometry gating on live CD3^+^CD4^+^ T cells. Average percentage of cells in the gates for all mice in each group is indicated. **(C)** Percentages of CD4 T cells positive for Tbet, GATA3, RORγt, or FOXP3 transcription factors, analyzed by intracellular flow cytometry gating live CD3^+^CD4^+^ T cells. Positive gates for each channel were set with an FMO control. Bars represent means ± SEM; statistical analyses with one-way ANOVA and Sidak’s post-hoc test; * *p*<0.05, ** *p*<0.01, *** *p*<0.001; n=5-6 mice per group. **(D)** Representative flow cytometry plots, showing the FOXP3^+^ Treg cells among live CD3^+^CD4^+^ T cells, in co-cultures with DCs from SKG-RA and control SKG-PBS mice. The average percentage of cells in the gate for all mice in each group is indicated.

We further analyzed the capacity of RA-SKG DCs to induce the expression of T-helper cell transcription factors Tbet, GATA3, RORγt, and FOXP3, corresponding to Th1, Th2, Th17, and Treg lineages, respectively. The reduced induction of Tbet in CD4 T cells was observed in co-cultures with RA-SKG DCs under all the stimulation conditions, while reduced induction of GATA3 was observed under β-glucan stimulation only, and the induction of RORγt was not significantly affected (Figure 6C). Importantly, RA-SKG DCs were impaired in the induction of Treg transcription factor FOXP3 across all the stimulation conditions (Figure 6C-D), consistent with their previously established impaired capacity for IL-10 induction (Figure 6A-B, Figure S12A).

T cells co-cultured with RA-SKG DCs also showed reduced induction of the activation marker CD44, increased levels of the naïve T cell marker CD62L, but unchanged induction of the activation marker CD69 and exhaustion marker CTLA4, as compared to the T cells in co-cultures with control DCs (Figure S12B-E). T cells co-cultured with RA-SKG DCs also showed a reduced induction of the proliferation marker Ki-67 (Figure S12D-E), although T cell proliferation as measured with a dye-dilution assay was variable across the biological replicates and overall not significantly affected (data not shown). We conclude that the cell-intrinsic impact of arthritis on DC differentiation results in the production of DCs with profoundly altered capacity for T cell activation. In particular, the reduced capacity of DCs derived from arthritis afflicted mice to induce FOXP3^+^ and IL-10^+^ CD4 T cells may contribute to the pathogenesis of arthritis.

## Discussion

In summary, in the current work we establish that the expansion of HSPCs and the induction of emergency myelopoiesis in murine rheumatoid arthritis is associated with enhanced DC production and significantly altered DC capacity for costimulatory and checkpoint marker expression, cytokine production, and T cell activation. Such effects are cell intrinsic and can persist independently of the inflammatory disease milieu. This suggests an induction of trained immunity that affects DC development and function and could contribute to rheumatoid arthritis disease persistence and progression.

Trained immunity, involving long-term changes in HSPC transcriptional, metabolic, and epigenetic profiles (13, 14) and resulting in the downstream remodeling of innate immune cell differentiation and function, has become well established in vaccination and infection models (11, 12), and there is a growing interest in its role in chronic inflammatory pathologies (18, 19). While most studies focus on monocytes and macrophages as the major mediators of trained immunity, in recent years similar mechanisms have been implicated for other innate immune cell lineages, including neutrophils (66, 67, 68, 69), osteoclasts (18, 70), and mast cells (71). Modulation of DC function through trained immunity mechanisms is indicated in two recent studies in murine models, namely with *C. neoformans* strain H99γ immunization (37) and intradermal inoculation with cholera toxin B subunit (38). In both models, enhanced DC responses to recall stimulation were reported, correlating with protection against virulent *C. neoformans* infection (37) and enhanced antitumor immunity in a melanoma model (38), respectively. Importantly, our current work for the first time expands such analyses of DC trained immunity to rheumatoid arthritis, representing a common chronic autoimmune and inflammatory disease and a major burden on the healthcare system.

The precise impact of altered DC differentiation and function on arthritis disease mechanisms remains to be further investigated. Our initial analyses suggested an impaired capacity of both macrophages and DCs derived from the arthritis afflicted mice to respond to microbial stimulation, based on reduced expression of costimulatory receptors and inflammatory cytokines and the enhanced expression of PD-L1 and IL-10. Subsequent global transcriptional analyses confirmed these effects, however also demonstrated a strong induction of many other inflammatory and immunogenic gene signatures in the RA-SKG DCs. It is notable that in previous studies in murine infection and vaccination models IL-1β was required for effective induction of trained immunity (16, 72, 73, 74), while type-I IFN responses were associated with a failure to induce protective trained immunity (17). The downregulation of IL-1β production and the strong transcriptional induction of type-I IFN response in RA-SKG DCs in the current study therefore indicates the distinct immunological signatures and mechanisms of trained immunity in rheumatoid arthritis versus these classical models.

Nevertheless, the notable dysregulation of major immune effectors, like IL-12, IL-10, PD-L1 and others, in both DCs and macrophages from arthritis afflicted mice, demonstrated in our current study, may indicate a dampened antimicrobial activity and an impaired capacity to mount a protective immune response against certain infectious challenges. Indeed, the increased incidence and severity of infections in patients with rheumatoid arthritis is well documented, and is at least in part independent from immunosuppressive therapies (75, 76, 77). Our studies suggest a potential role for immunosuppressive trained immunity induced in response to chronic inflammation in such clinical outcomes, and future studies with human patients may address the underlying mechanisms.

DCs are the most potent antigen presenting cells with unique capacity for activation of naïve T cells and priming of antigen specific immune responses (26, 27). With the central role of autoreactive T cells in driving arthritis pathology in the SKG model and in human patients, it will be essential to establish the impact of DC trained immunity, as described in our current work, on both antigen specific and heterologous T cell activation. Crosstalk between trained immunity and the adaptive immune system was indicated in previous studies (78, 79), although without implicating altered DC development and function. For example, trained immunity induced through BCG vaccination was shown to cross-protect against influenza A infection in mouse models by inducing an increase in memory CD4 T cells in circulation and in the lungs, with IFNγ derived from T cell activating the antimicrobial functions of alveolar macrophages (80). Similarly, induction but not maintenance of memory-like alveolar macrophages following respiratory viral infections in mice was shown to be dependent on IFNγ production by CD8 T cells (81). In the current study, DCs derived ex vivo from the bone marrow of arthritis afflicted mice had an altered capacity for CD4 T cell activation, resulting in reduced induction of T cell activation markers, cytokines, and Th-lineage transcription factors, but retained induction of T cell proliferation. Overall, this indicates that the alterations in DC functions through the trained immunity mechanisms may result in downstream changes in T cell mediated adaptive immune responses.

Beyond their immunogenic functions, the capacity of DCs to induce distinct immune response polarizations needs to be considered when assessing their role in disease pathologies (82). Arthritis onset and progression in SKG mice is linked to the expansion of autoreactive Th17 cells (83, 84), and Th17-polarized responses are also implicated in disease pathogenesis in human rheumatic diseases (85, 86, 87). It is therefore notable that in the current study DCs from arthritis afflicted mice induced lower levels of Tbet^+^ but not RORγt^+^ CD4 T cells, although the induction of both IFNγ and IL-17A was reduced at least under some of the conditions tested. Nevertheless, the reduced induction of IL-10^+^ and FoxP3^+^ CD4 T cells likely indicates a more immunogenic state of RA-SKG DCs and could contribute to disease pathogenesis in vivo.

As well as expanding the fundamental knowledge of immune dysfunction in rheumatoid arthritis, the current study may advance the understanding of the mechanisms of action of existing therapies and potentially suggest novel targets for therapeutic intervention. Understanding the mechanisms of trained immunity is particularly important with the growing application of autologous HSPC transplantation as a therapy for severe autoimmune and rheumatic diseases (88, 89, 90). It will be essential to establish if persisting functional changes in HSPCs and innate trained immunity favors disease remission or relapse following autologous HSPC transplant, and to develop approaches to either facilitate or mitigate such effects.

## Ethics Statement

Mouse experiments were in accordance with guidelines of the Canadian Council on Animal Care and protocols MCGL-7932 and 6029 approved by the McGill Animal Care Committee. Studies with human subjects were approved by the McGill University Health Center Ethics Review Board (ERB protocol number MP-37-2021-7495), and written consent was obtained from all the participants.

## Data Availability Statement

RNA-seq data will be available from the National Center for Biotechnology Information GEO database upon the publication of the paper. All other data are available from the corresponding authors upon request.

## Author Contributions Statement

Data were acquired by LTT, RRJ, BP, and LD with the support from JK, MM, and YL, and with the supervision of AN, DL, SMV, DM, MY, IC, and CK. The disease models were established by MY, SMV, and DM. Experiments were designed by LTT, RRJ, AN, SMV, DL, DM, MY, and CK. The data were analyzed and prepared for publication by LTT, RRJ, MM, AN, and DL. The manuscript was written and edited by AN, LTT, RRJ, DL, IC, CK, SVM, and reviewed by other co-authors.

## Acknowledgments

We thank Patricia D’Arcy for mouse colony management, disease induction, and monitoring. We thank McGill Comparative Medicine and Animal Resources Centre (CMARC) for mouse husbandry and veterinary services. We thank Sonia Léger-Thériault, Clinical Research Coordinator-Nurse at RI-MUHC, for patient recruitment and blood sample collection. Flow cytometry was performed at McGill Life Sciences Complex, and we thank facility managers Julien Leconte and Camille Stegen. RNA-seq was performed with the Molecular Biology and Functional Genomics Facility of the Institut de Recherches Cliniques de Montréal (IRCM), and we thank Sarah Boissel and other staff of the facility. We thank the Digital Research Alliance of Canada and Calcul Québec for the resources for bioinformatics data analyses. We thank the CMARC Diagnostics Laboratory for hematology analyses of mouse blood.

