## Supplemental for "Trained Immunity Affecting Dendritic Cell Differentiation and Function in Rheumatoid Arthritis"

### Supplemental Tables

**Supplemental Table S1. Antibodies for flow cytometry analyses of mouse tissues.**

| Target | Fluorophore | Company | Clone | Catalogue |
|---|---|---|---|---|
| CD3 | Biotin | Biolegend | 17A2 | 100244 |
| B220 | Biotin | Biolegend | RA3-6B2 | 103203 |
| TER119 | Biotin | Biolegend | TER-119 | 116204 |
| Streptavidin | Brilliant Violet 785 | Biolegend | NA | 405249 |
| cKIT | Brilliant Violet 650 | Biolegend | ACK2 | 135125 |
| SCA1 | Brilliant Ultraviolet 395 | BD Biosciences | D7 | 563990 |
| FLT3 | PE | eBioscience<br>Thermo Fisher Scientific | A2F10 | 12-1351-83 |
| CD34 | Brilliant Violet 421 | Biolegend | SA376A4 | 152208 |
| CD150 | PE-Cy7 | Biolegend | TC15-12F12.2 | 115913 |
| CD48 | PerCP-Cy5.5 | Biolegend | HM48-1 | 103422 |
| CD16/CD32 | Brilliant Ultraviolet 737 | BD Biosciences | 2.4G2 | 612783 |
| TREM1 | eFlour660 | eBioscience<br>Thermo Fisher Scientific | TR3MBL1 | 50-3541-80 |
| Dectin1 | FITC | eBioscience<br>Thermo Fisher Scientific | 2A11 | MA5-16479 |
| TLR2 | APC | Biolegend | QA16A01 | 153006 |
| CD14 | FITC | Biolegend | Sa14-2 | 123308 |
| CD45 | Brilliant Ultraviolet 395 | BD Biosciences | 30-F11 | 564279 |
| CD11b | eFlour450 | eBioscience<br>Thermo Fisher Scientific | M170 | 48-0112-82 |
| Ly6G | APC-Cy7 | Biolegend | 1A8 | 127624 |
| Ly6C | PerCP-Cy5.5 | Biolegend | HK1.4 | 128012 |
| CD11c | Brilliant Ultraviolet 737 | BD Biosciences | HL3 | 612796 |
| F4/80 | Brilliant Violet 785 | Biolegend | BM8 | 123141 |
| MHCII | Brilliant Violet 650 | Biolegend | M5/114 | 107641 |
| PD-L1 | PE-Cy7 | Biolegend | 10F.9G2 | 124314 |
| SiglecF | PE | Biolegend | S17007L | 155506 |
| CD86 | APC | Biolegend | GL1 | 105012 |
| CD40 | FITC | Biolegend | HM40-3 | 102906 |
| Ly6G | Biotin | Biolegend | 1A8 | 127604 |
| cKIT | PE-Cy7 | Biolegend | 2B8 | 105814 |
| SCA1 | APC | eBioscience<br>Thermo Fisher Scientific | D7 | 17-5981-83 |
| CD16/CD32 | FITC | eBioscience<br>Thermo Fisher Scientific | 93 | 11-0161-85 |
| CD172α | PerCP-Cy5.5 | Biolegend | P84 | 144010 |
| CD45.1 | Brilliant Ultraviolet 395 | BD Biosciences | A20 | 565212 |
| CD45.2 | PE-Cy7 | BioLegend | 104 | 109830 |

**Supplemental Table S2. Antibodies for ex vivo analyses of bone marrow derived macrophages.**

| Target | Fluorophore | Company | Clone | Catalogue |
|---|---|---|---|---|
| CD11b | eFlour450 | eBioscience Thermofisher Scientific | M170 | 48-0112-82 |
| F4/80 | Brilliant Violet 785 | BioLegend | BM8 | 123141 |
| PD-L1 | PE-Cy7 | BioLegend | 10F.9G2 | 124314 |
| MHCII | Brilliant Violet 650 | BioLegend | M5/114.15.2 | 107641 |
| CD80 | PE | eBioscience Thermofisher Scientific | 16-10A1 | 12-0801-82 |
| CD86 | APC | BioLegend | GL1 | 105012 |
| CD40 | FITC | BioLegend | HM40-3 | 102906 |
| PD-L2 | PerCPCy5.5 | BioLegend | TY25 | 107218 |

| Target | Fluorophore | Company | Clone | Catalogue |
|---|---|---|---|---|
| CD11b | eFlour450 | eBioscience Thermofisher Scientific | M170 | 48-0112-82 |
| F4/80 | Brilliant Violet 785 | BioLegend | BM8 | 123141 |
| *iNOS | PE | BD Biosciences | W16030C | 696806 |

* following processing with the Transcription Factor Staining Buffer Set (eBioscience #00-5523-00)

| Target | Fluorophore | Company | Clone | Catalogue |
|---|---|---|---|---|
| CD11b | eFlour450 | eBioscience Thermofisher Scientific | M170 | 48-0112-82 |
| F4/80 | Brilliant Violet 785 | BioLegend | BM8 | 123141 |
| *IL-6 | APC | BioLegend | MP5-20F3 | 504508 |
| *TNFα | FITC | BioLegend | MP6-XT22 | 506304 |

* following processing with the Fixation/Permeabilization Solution Kit (BD Biosciences #554714)

**Supplemental Table S3. Antibodies for ex vivo analyses of bone marrow derived dendritic cells.**

| Target | Fluorophore | Company | Clone | Catalogue |
|---|---|---|---|---|
| CD11b | eFluor 450 | eBioscience | m1/70 | 48-0112-82 |
| CD11c | Brilliant Ultraviolet 737 | eBioscience | N418 | 367-0114-82 |
| CD40 | FITC | BioLegend | HM40-3 | 102906 |
| CD80 | PE | eBioscience | 16-10A1 | 12-0801-82 |
| CD86 | APC | BioLegend | GL-1 | 105012 |
| F4/80 | BV785 | BioLegend | BM8 | 123141 |
| PD-L1 | PE-Cy7 | BioLegend | 10F.9G2 | 124314 |
| PD-L2 | PerCP-Cy5.5 | BioLegend | TY25 | 107218 |
| MHCII | BV650 | BioLegend | M5/114.15.2 | 107641 |

**Supplemental Table S4. Antibodies for ex vivo analyses of murine T cell activation.**

| Target | Fluorophore | Company | Clone | Catalogue |
|---|---|---|---|---|
| IFNγ | PE | BioLegend | W18272D | 163503 |
| IL-2 | Brilliant Violet 421 | BioLegend | JES6-5H4 | 503825 |
| IL-4 | Alexa Fluor 647 | BioLegend | 11B11 | 504112 |
| IL-10 | PE-Cy7 | BioLegend | JES5-16E3 | 505025 |
| T-bet | PE | BioLegend | 4B10 | 644809 |
| GATA3 | Alexa Fluor 647 | BioLegend | 16E10A23 | 653809 |
| RORγt | Brilliant Violet 421 | BD Biosciences | Q31-378 | 562894 |
| FoxP3 | FITC | ThermoFisher | FJK-16s | 11-5773-82 |
| Ki-67 | Brilliant Violet 650 | BioLegend | 11F6 | 151215 |
| CD3 | Brilliant Ultraviolet 395 | BD Biosciences | 145-2C11 | 563565 |
| CD4 | PerCP-Cy5.5 | Biolegend | RM4-4 | 116012 |

**Supplemental Table S5. Demographic characteristics of study participants.**
Abbreviations: DM: diabetes mellitus, HTN: hypertension, HC: hypercholesterolemia.

| | Controls (n=4) | RA (n=4) |
|---|---|---|
| Age (mean ± SD) | 47.2 +/- 7 | 43.5 +/- 6.1 |
| Sex (F) | 3 | 3 |
| Comorbidities | | |
| Smoking (Y) | 1 | 1 |
| DM (Y) | 1 | 0 |
| HTN | 1 | 0 |
| HC | 1 | 0 |
| RA Medications | | |
| Methotrexate | n/a | 2 |
| Upadacitinib/Leflunomide) | n/a | 1 |
| Prednisone (3mg) | n/a | 1 |

**Supplemental Table S6. Antibodies for the analyses of human monocyte derived DCs.**

| | | |
|---|---|---|
| CD14 | BV650 | BioLegend 301835 Brilliant Violet 650 anti-human CD14 |
| CD16 | BUV737 | BD Biosciences 612786 BUV737 anti-human CD16 |
| CD11c | BV421 | BioLegend 337226 Brilliant Violet 421 anti-human CD11c |
| DC-SIGN | PerCP-Cy5.5 | BioLegend 330109 PerCP/Cyanine5.5 anti-human CD209 (DC-SIGN) |
| HLA-DR | PE-Cy7 | BioLegend 307616 PE/Cy7 anti-human HLA-DR |
| CD86 | APC | BioLegend 374207 APC anti-human CD86 |
| CD40 | FITC | BioLegend 334305 FITC anti-human CD40 Antibody |
| PD-L1 | PE | BioLegend 329705 PE anti-human CD274 (B7-H1, PD-L1) |

**Supplementary Table S7. RNA-seq transcriptional analyses of BMDCs derived from lineage depleted bone marrow of arthritis mice; (*to be attached as a separate excel file at the time of publication*).** **(A)** Full list of genes expressed in BMDCs derived from SKG mice under PBS-NS (non-stimulated), PBS-BDG (β-Glucan), PBS-PIC (poly(I:C)), Zym-NS, Zym-BDG, and Zym-PIC conditions. Information provided for each gene includes: gene name and counts per million (CPM). **(B)** List of genes differentially expressed in non-stimulated BMDCs (NS) derived from mice with active arthritis disease (Zym-NS) relative to BMDCs derived from control mice (PBS-NS). **(C)** List of genes differentially expressed in BDG-stimulated BMDCs derived from mice with active arthritis disease (Zym-BDG) relative to BMDCs derived from control mice (PBS-BDG). **(D)** List of genes differentially expressed in PIC-stimulated BMDCs derived from mice with active arthritis disease (Zym-PIC) relative to BMDCs derived from control mice (PBS-PIC). **(B-D)** Differential gene expression analyzed at fold change (FC) $\geq$ 2.0 and false discovery rate (FDR) $\leq$ 0.001; related to Figure 5. Information provided for each gene includes: gene name, fold change, normalized counts per million (CPM), *p*-value, and false discovery rate (FDR). **(E-G)** Gene set enrichment analyses (GSEA) showing select enriched biological process (BP) terms of differentially expressed genes from each of the three comparisons (Zym-NS vs PBS-NS, Zym-BDG vs PBS-BDG, and Zym-PIC vs PBS-PIC). In the normalized enrichment scores (NES) column, positive values indicate upregulation and negative values indicate downregulation in the comparisons.

#### SUPPLEMENTAL FIGURES (WITH LEGENDS)

**Figure S1. Induction of emergency myelopoiesis in murine arthritis. (A)** Hematology analysis of the blood of RA-SKG and control SKG mice; MCV – mean corpuscular volume; MCHC – mean corpuscular hemoglobin concentration. Bars represent means ± SEM; statistical analyses with unpaired *t*-tests; * $p<0.05$, ** $p<0.01$, *** $p<0.001$, ns – not significant; n=6-8 mice per group, consolidated from two independent experiments. **(B)** Monocytes and macrophage numbers in the bone marrow and spleen of RA-SKG and control SKG mice. Cells are gated as live $CD45^{+}CD11b^{+}$ $Ly6G^{-}SiglecF^{-}$, followed by $CD64^{-}F4/80^{-}Ly6C^{+}$ for monocytes, $CD64^{+}F4/80^{+}Ly6C^{+}$ for monocyte-derived macrophages, and $CD64^{+}F4/80^{+}Ly6C^{-}$ for tissue-resident macrophages. Bars represent means ± SEM; statistical analyses with unpaired *t*-tests; * $p<0.05$, ** $p<0.01$, ns – not significant; n=9 mice per group across two independent experiments. **(C)** Representative flow cytometry plots, gated on live $CD45^{+}CD11b^{+}Ly6G^{-}SiglecF^{-}$ cells, and showing monocytes, monocyte-derived macrophages, and tissue resident macrophages in the spleen of RA-SKG and control mice. Average percentage of cells in each gate for all mice in each group is shown. **(D)** Expression of activation and checkpoint markers CD40, CD86, MHCII and PD-L1 on monocytes in the bone marrow and spleen of RA-SKG and control SKG mice. Bars represent means ± SEM; statistical analyses with unpaired *t*-tests; * $p<0.05$, ** $p<0.01$, *** $p<0.001$, **** $p<0.0001$, ns – not significant; n=5 mice per group. **(E)** Representative histograms showing the expression of activation maker CD40 and checkpoint marker PD-L1 on monocytes in the bone marrow and spleen of RA-SKG and control mice. Monocytes are gated as live $CD45^{+}CD11b^{+}Ly6G^{-}SiglecF^{-}F4/80^{-}$ $Ly6C^{+}$ cells; MFI – mean fluorescence intensity.

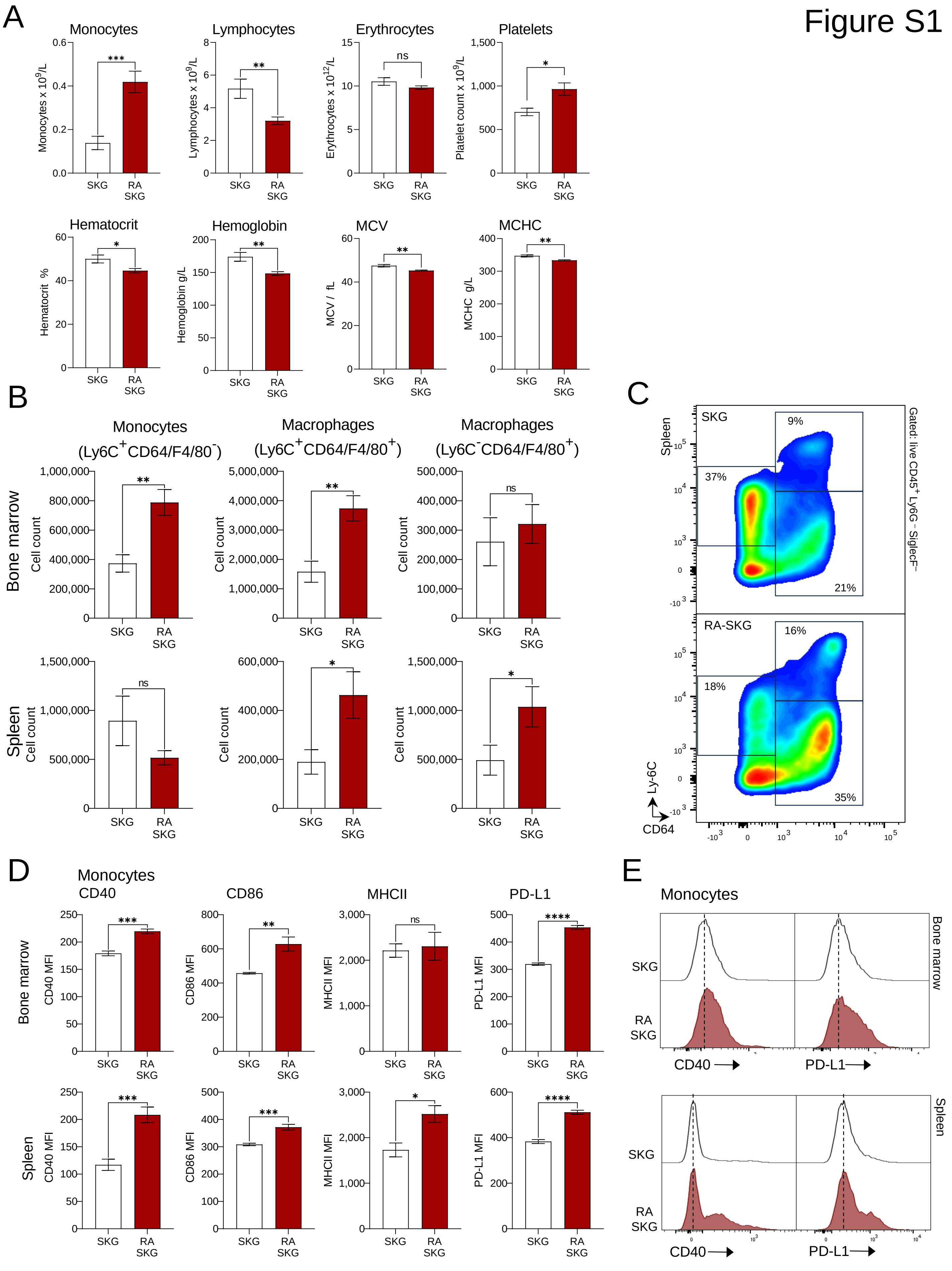
Figure S1
A
Monocytes
Lymphocytes
Erythrocytes
Platelets
Monocytes x 10^9/L
Lymphocytes x 10^9/L
Erythrocytes x 10^12/L
Platelet count x 10^9/L
Hematocrit
Hemoglobin
MCV
MCHC
Hematocrit %
Hemoglobin g/L
MCV / fL
MCHC g/L
SKG
RA SKG
B
Monocytes (Ly6C+CD64/F4/80-)
Macrophages (Ly6C+CD64/F4/80+)
Macrophages (Ly6C-CD64/F4/80+)
Bone marrow
Spleen
Cell count
C
SKG
RA-SKG
9%
37%
21%
16%
18%
35%
Gated: live CD45+ Ly6G- SiglecF-
Ly-6C
CD64
D
Monocytes
CD40
CD86
MHCII
PD-L1
CD40 MFI
CD86 MFI
MHCII MFI
PD-L1 MFI
E
Monocytes
CD40
PD-L1

**Figure S2. Markers and gating strategies for the analyses of monocytes, macrophages, and dendritic cells in murine tissues. (A)** Flow cytometry analysis of monocytes, macrophages, and other myeloid cells in murine tissues. The cells were gated as live $CD45^{+}CD11b^{+}Ly6G^{-}SiglecF^{-}$, followed by $CD64^{-}F4/80^{-}$ $Ly6C^{+}$ for monocytes, $CD64^{+}F4/80^{+}Ly6C^{+}$ for monocyte-derived macrophages, and $CD64^{+}F4/80^{+}Ly6C^{-}$ for tissue-resident macrophages. **(B)** Flow cytometry analysis of DCs in murine tissues. DCs were gated as live $CD45^{+}Lin^{-}F4/80^{-}CD64^{-}CD11c^{+}MHCII^{+}$ cells, followed by $B220^{-}PDCA1^{-}XCR1^{+}SIRP\alpha^{-}$ for cDC1, $B220^{-}PDCA1^{-}XCR1^{-}SIRP\alpha^{+}CLEC12a^{-}$ for cDC2a, $B220^{-}PDCA1^{-}XCR1^{-}SIRP\alpha^{+}CLEC12a^{+}$ for cDC2b, and $B220^{+}PDCA1^{+}$ for pDCs. Gating shown is for a sample of mouse spleen, with bone marrow using the same overall gating hierarchy with adjustments to gate positions.

### Figure S2

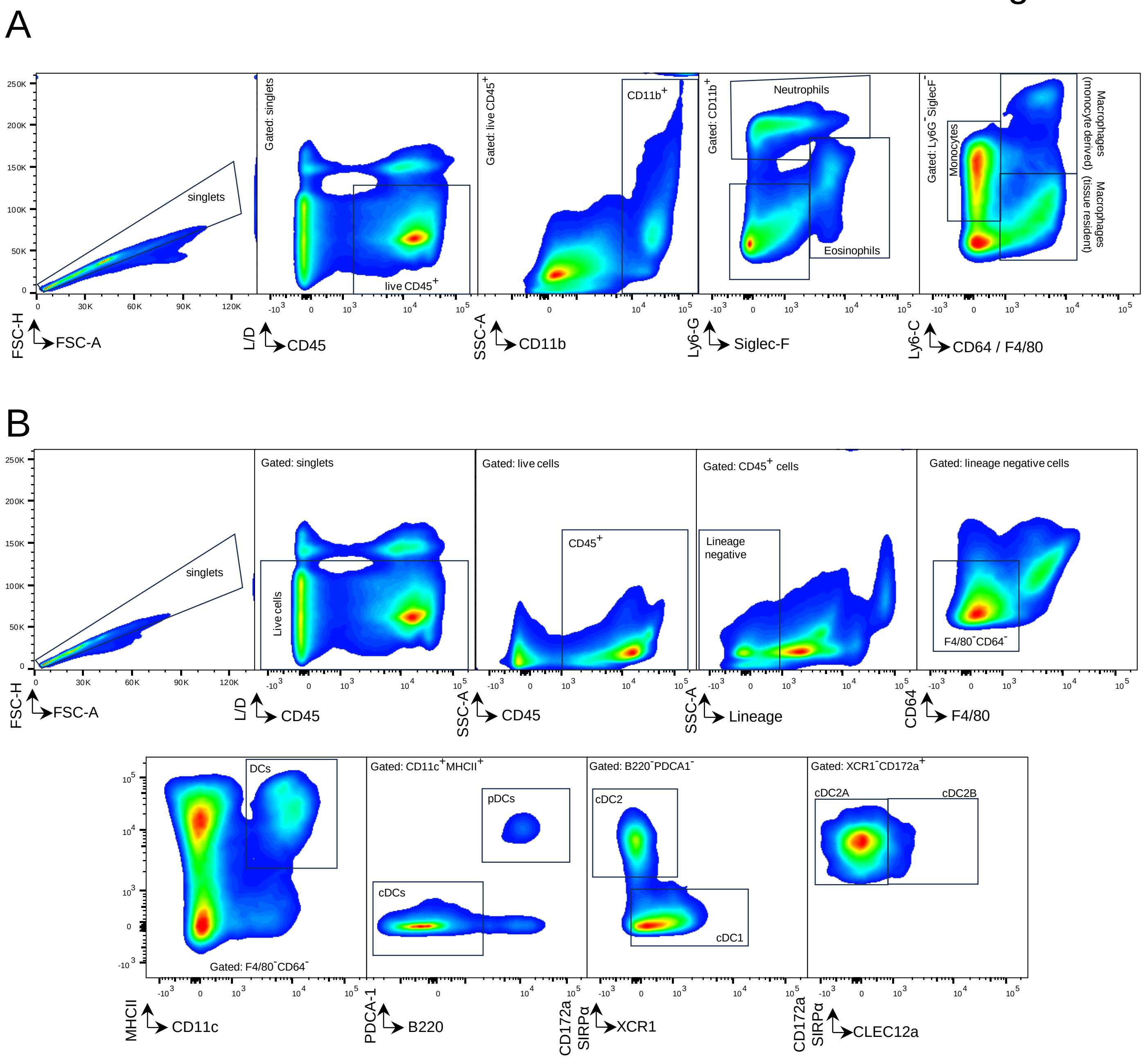
A
singlets
Gated: singlets
live CD45+
Gated: live CD45+
CD11b+
Gated: CD11b+
Neutrophils
Eosinophils
Gated: Ly6G-SiglecF-
Monocytes
Macrophages (monocyte derived)
Macrophages (tissue resident)
FSC-H FSC-A
L/D CD45
SSC-A CD11b
Ly6-G Siglec-F
Ly6-C CD64 / F4/80
B
singlets
Gated: singlets
Live cells
Gated: live cells
CD45+
Gated: CD45+ cells
Lineage negative
Gated: lineage negative cells
F4/80-CD64-
FSC-H FSC-A
L/D CD45
SSC-A CD45
SSC-A Lineage
CD64 F4/80
DCs
Gated: F4/80-CD64-
Gated: CD11c+MHCII+
pDCs
cDCs
Gated: B220-PDCA1-
cDC2
cDC1
Gated: XCR1-CD172a+
cDC2A
cDC2B
MHCII CD11c
PDCA-1 B220
CD172a SIRPα XCR1
CD172a SIRPα CLEC12a

**Figure S3. Bone marrow myeloid colony forming units and the expression of pattern recognition receptors on hematopoietic progenitor cells in arthritis afflicted SKG mice. (A)** Myeloid colony forming units (CFUs) in the bone marrow of SKG-RA mice, with the counts normalized for each experiment to the average levels of myeloid CFUs in the control SKG group. **(B)** Expression of TLR2, CD14, TREM1, and Dectin1 on common myeloid progenitors (CMPs), granulocyte monocyte progenitors (GMP), megakaryocyte progenitors (MkP), and megakaryocyte erythroid progenitors (MEPs), quantified as mean fluorescence intensity (MFI). **(C)** Representative histograms showing the expression of TLR2, CD14, TREM1, and Dectin1 on CMPs in arthritis afflicted and control SKG mice. Cells were gated as $Lin^{-}cKit^{+}Sca1^{-}$ followed by $CD34^{+}CD16/32^{-}$ for common myeloid progenitors (CMP), $CD34^{+}CD16/32^{+}$ for granulocyte monocyte progenitors (GMP), $CD34^{-}CD16/32^{-}$ for megakaryocyte erythroid progenitor (MEP), and $CD41^{+}CD150^{+}$ for megakaryocyte progenitors (MkP). Bars represent means ± SEM; data is from 5-15 mice per group consolidated from 2-3 independent experiments; statistical analyses with Mann-Whitney test; * $p$<0.05, ** $p$<0.01, *** $p$<0.001, ns – not significant; MFI – mean or median fluorescence intensity. Isotype controls were used throughout to confirm positive staining.

Figure S3

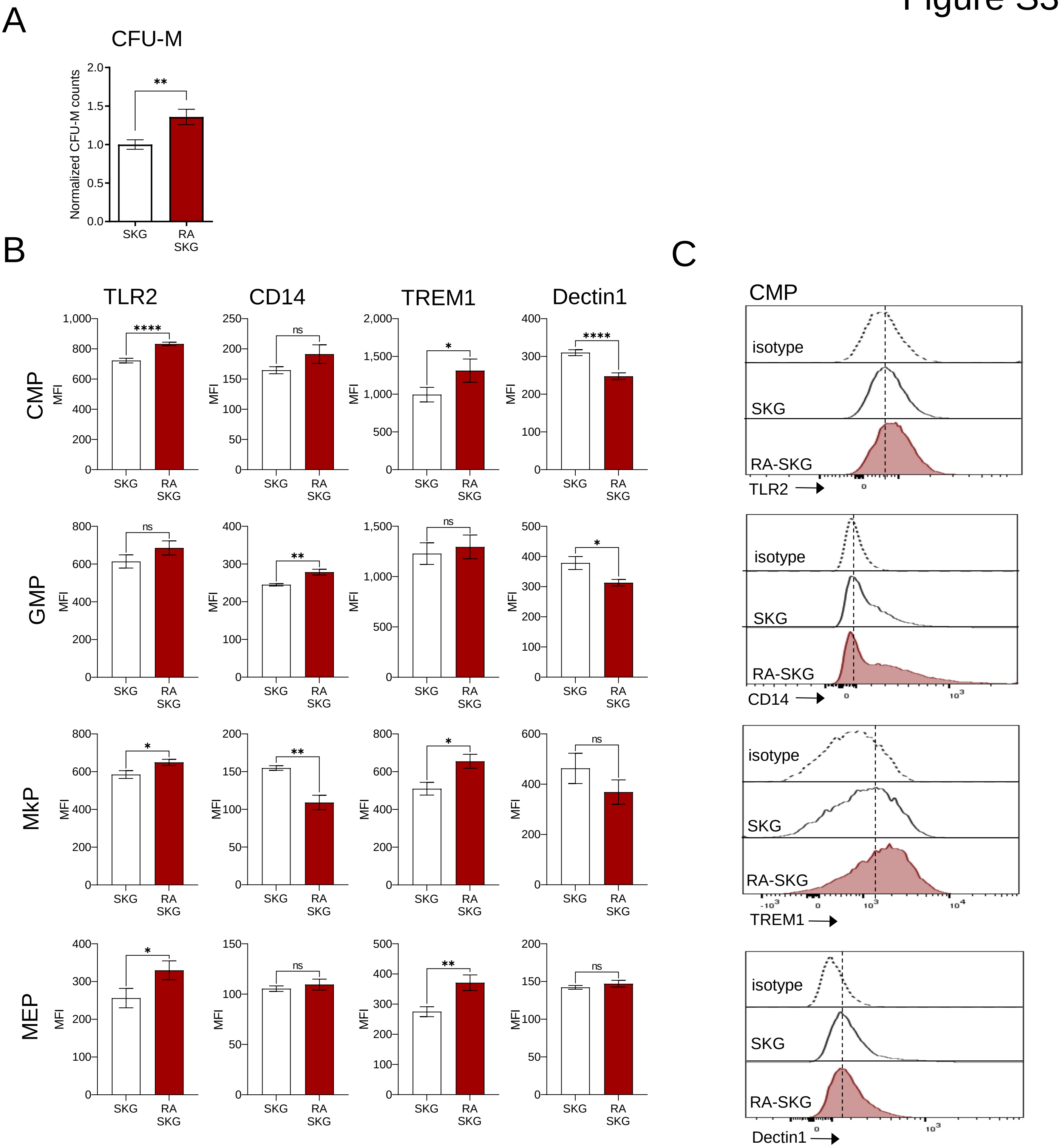

A
CFU-M
Normalized CFU-M counts
**
SKG
RA SKG
B
TLR2
CD14
TREM1
Dectin1
CMP
GMP
MkP
MEP
MFI
****
ns
*
**
SKG
RA SKG
C
CMP
isotype
SKG
RA-SKG
TLR2
CD14
TREM1
Dectin1

**Figure S4. Altered functional profile of macrophages derived *ex vivo* from bone marrow of mice with active arthritis disease.** BMDMs were derived *ex vivo* from the bone marrow of SKG-RA and control SKG-PBS mice, and stimulated with LPS 10 ng/mL, β-glucan 5 μg/mL, or poly(I:C) 20 μg/mL for 12 hours. **(A-B)** Analyses of IL-6 and TNFα production by the BMDMs through intracellular flow cytometry, presented as the percentages of cells positive for the cytokines, pre-gating on live $CD11b^+$ $F4/80^+$ cells. **(C-E)** Analyses of iNOS levels in BMDMs through intracellular flow cytometry, including **(C)** the percentages of iNOS positive cells, **(D)** the mean fluorescence intensity (MFI) of iNOS staining, and **(E)** representative flow cytometry plots showing iNOS staining, all pre-gated on live $CD11b^+$ $F4/80^+$ cells; percentages of cells in the positive gates are shown as mean ± S.D. for each group. Bars represent means ± SEM; n=4 mice per group; statistical analyses with one-way ANOVA and Sidak's post-hoc test; * $p<0.05$, ** $p<0.01$, *** $p<0.001$.

### Figure S4

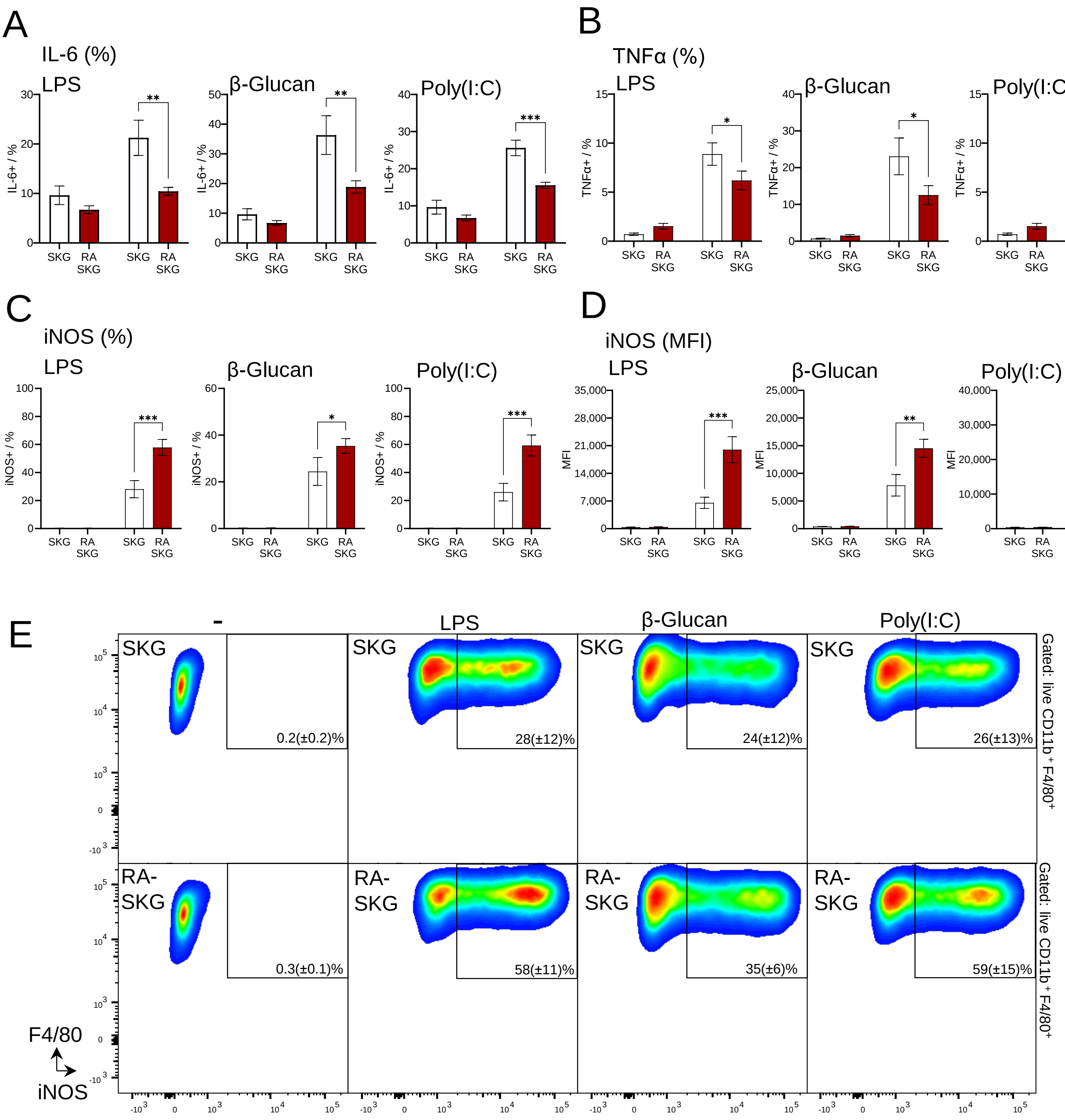

A
IL-6 (%)
LPS
β-Glucan
Poly(I:C)
IL-6+ / %
SKG
RA SKG
**
***
B
TNFα (%)
LPS
β-Glucan
Poly(I:C)
TNFα+ / %
*
ns
C
iNOS (%)
iNOS+ / %
D
iNOS (MFI)
MFI
E
-
SKG
RA-SKG
0.2(±0.2)%
28(±12)%
24(±12)%
26(±13)%
0.3(±0.1)%
58(±11)%
35(±6)%
59(±15)%
Gated: live CD11b+ F4/80+
F4/80
iNOS

**Figure S5. Further characterization of dendritic cell development in the SKG mouse model of rheumatoid arthritis (A-B)** Expansion on pre-DCs in the bone marrow of arthritis afflicted SKG mice. **(A)** Increased numbers of pre-DCs in the bone marrow of arthritis afflicted SKG mice. Cells were gated as live $Lin^{-}Sca1^{-}cKit^{+/low}CD115^{-}Flt3^{+}CD11c^{+}$. **(B)** Representative flow cytometry plots of mouse bone marrow pre-gated on live $Lin^{-}Sca1^{-}cKit^{+/low}CD115^{-}$ cells and showing the $Flt3^{+}CD11c^{+}$ pre-DC cell population. Percentage of cells in the pre-DC gate out of the overall live cells within the samples is indicated as mean ± S.D. for all the mice in each group. **(C-F)** Validation of BMDC production from lineage-depleted murine bone marrow cultures. **(C)** Yields of BMDCs, corresponding to non-adherent cells in the GM-CSF supplemented cultures of lineage depleted mouse bone marrow (day 8), quantified as cell counts on a hemocytometer. **(D)** Frequencies of BMDCs, quantified with flow cytometry as live $CD11b^{+}CD11c^{+}$ cells in the bone marrow cultures, including stimulation with LPS 10 ng/mL, β-glucan 5 μg/mL, or poly(I:C) 20 μg/mL for 16 hours. **(E)** Representative flow cytometry analyses of the BMDC cultures from SKG-RA and control SKG-PBS mice, gated on live cells, with the $CD11b^{+}CD11c^{+}$ BMDCs gate indicated. **(F)** Yields of live $CD11b^{+}CD11c^{+}$ BMDCs, calculated from the overall number of non-adherent cells and the frequency of live $CD11b^{+}CD11c^{+}$ BMDCs in the cultures. Bars represent means ± SEM; statistical analyses with unpaired *t*-test for two datasets or with one-way ANOVA and Sidak's post-hoc test; * $p<0.05$, ** $p<0.01$, *** $p<0.001$; n≥4 mice per group.

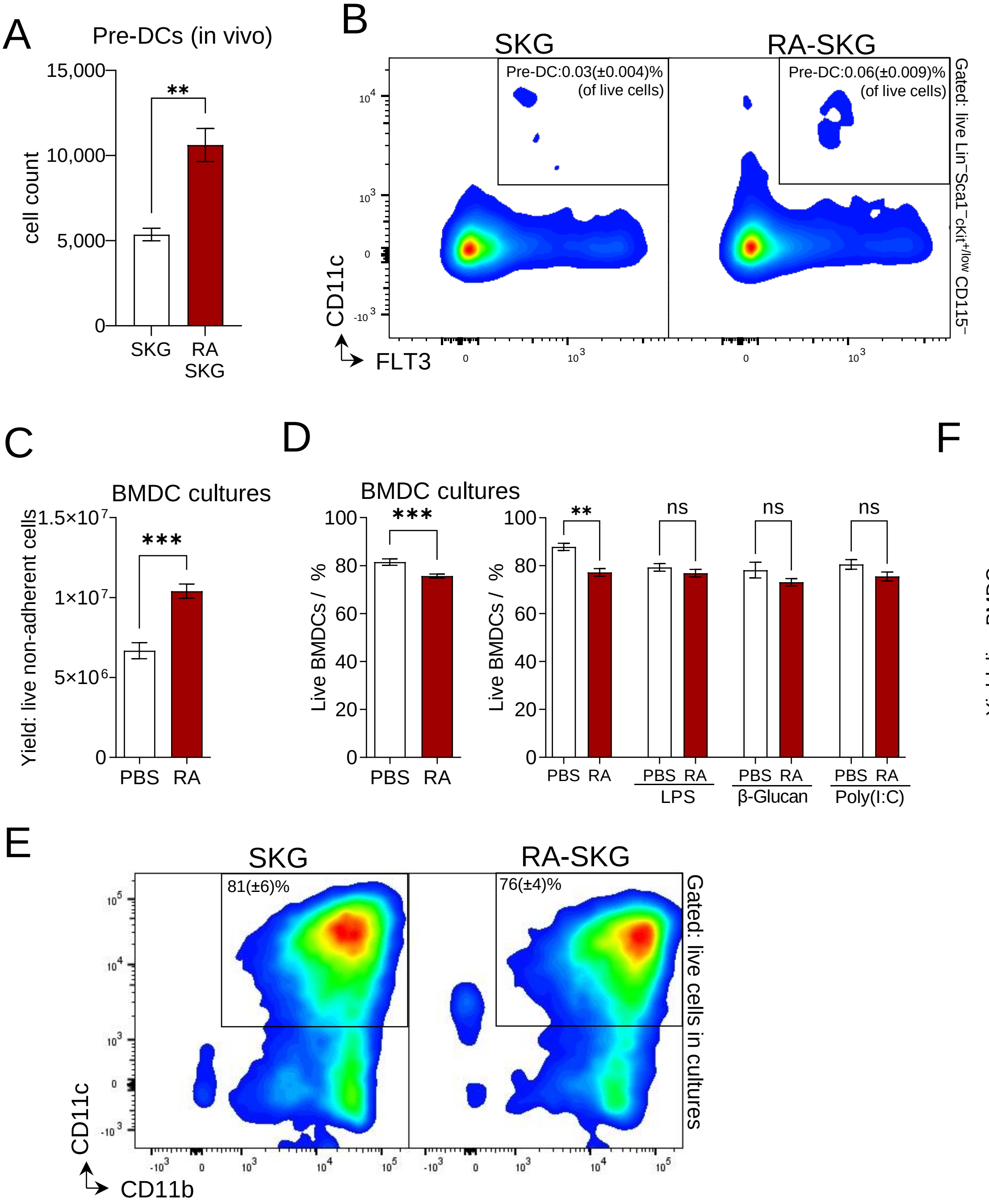
A
Pre-DCs (in vivo)
cell count
15,000
10,000
5,000
0
**
SKG
RA SKG
B
SKG
RA-SKG
Pre-DC:0.03(±0.004)% (of live cells)
Pre-DC:0.06(±0.009)% (of live cells)
Gated: live Lin⁻Sca1⁻cKit+/low CD115⁻
CD11c
FLT3
C
BMDC cultures
Yield: live non-adherent cells
1.5×10⁷
1×10⁷
5×10⁶
0
***
PBS
RA
D
BMDC cultures
Live BMDCs / %
100
80
60
40
20
0
***
PBS
RA
**
ns
ns
ns
LPS
β-Glucan
Poly(I:C)
F
BMDC cultures
Yield: live BMDCs
1×10⁷
8×10⁶
6×10⁶
4×10⁶
2×10⁶
0
**
PBS
RA
E
SKG
RA-SKG
81(±6)%
76(±4)%
Gated: live cells in cultures
CD11c
CD11b

**Figure S6. Altered functional profile of dendritic cells derived *ex vivo* from the bone marrow of mice with active arthritis disease: SKG arthritis model.** BMDCs were derived *ex vivo* from non-depleted bone marrow of SKG mice with arthritis disease, and subsequently stimulated with LPS 10 ng/mL, β-glucan 5 μg/mL, or poly(I:C) 20 μg/mL for 16 hours. **(A)** Expression of activation and checkpoint markers on BMDCs, analyzed by flow cytometry, pre-gating on live $CD11b^{+}CD11c^{+}$ cells; MFI – mean fluorescence intensity. Bars represent means ± SEM; statistical analyses with one-way ANOVA and Sidak's post-hoc test; * $p<0.05$, ** $p<0.01$, *** $p<0.001$; n=5 mice per group, and the results were reproduced in a second independent experiment. **(B)** Representative flow cytometry histograms showing the expression of activation and checkpoint markers on pre-stimulated BMDCs, gating on live $CD11b^{+}CD11c^{+}$ cells.

Figure S6

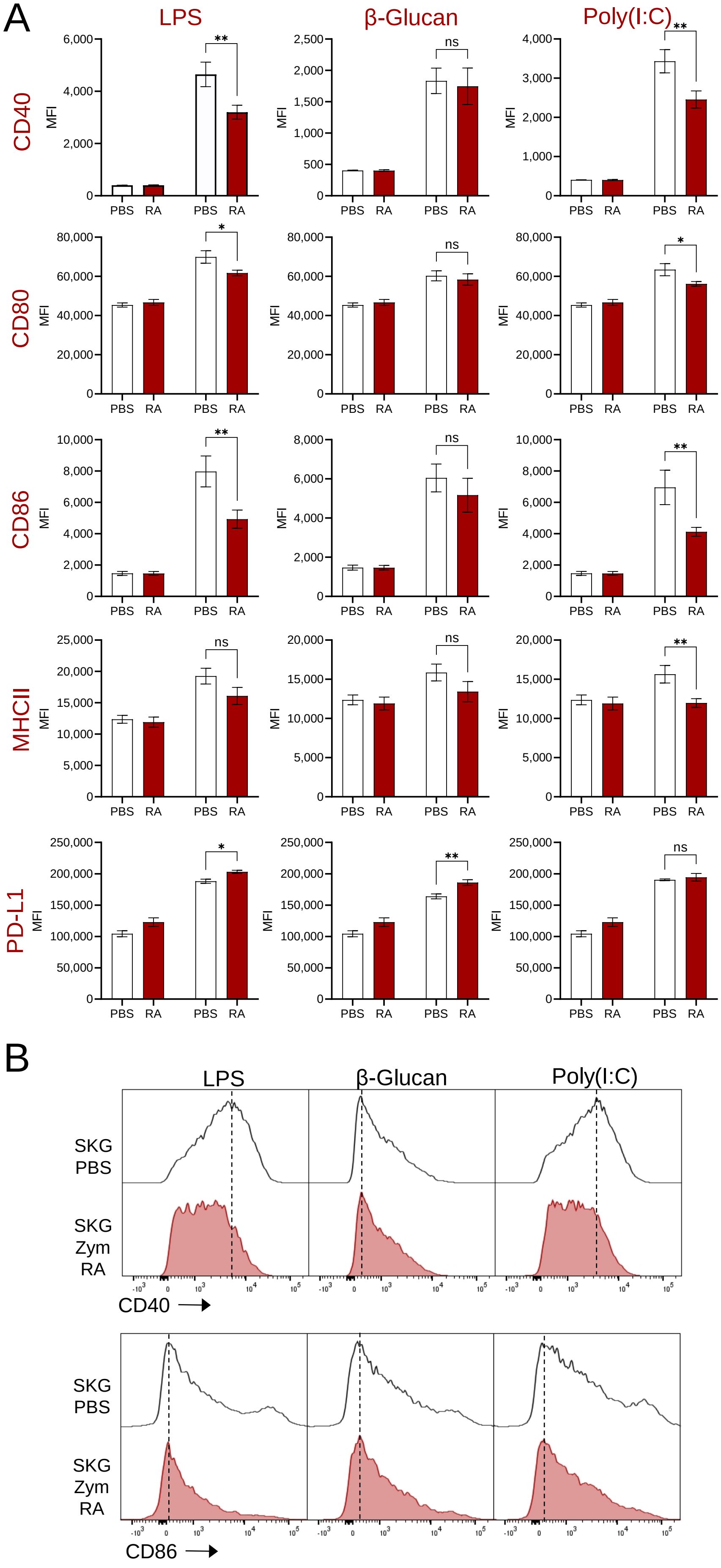
A
LPS
β-Glucan
Poly(I:C)
CD40
CD80
CD86
MHCII
PD-L1
MFI
PBS
RA
ns
*
**
B
LPS
β-Glucan
Poly(I:C)
SKG PBS
SKG Zym RA
CD40
CD86

**Figure S7. Altered functional profile of the *ex vivo* derived murine dendritic cells is the result of arthritis disease, and not zymosan treatment.** BMDCs were derived *ex vivo* from the bone marrow of wild type or SKG BALB/c mice, pre-treated with PBS or zymosan (5mg, Zym), 5.5 weeks earlier. Of the four mouse groups only SKG-Zym mice develop arthritis disease, while the others mount transient inflammatory response and subsequently remain healthy. BMDCs were stimulated with LPS 10 ng/mL, β-glucan 5 μg/mL, or poly(I:C) 20 μg/mL for 16 hours. **(A)** Expression of activation and checkpoint markers on BMDCs, analyzed by flow cytometry, pre-gating on live $CD11b^{+}CD11c^{+}$ cells; MFI – mean fluorescence intensity. Bars represent means ± SEM; statistical analyses with one-way ANOVA and Sidak's post-hoc test; * $p<0.05$, ** $p<0.01$, *** $p<0.001$; n=3-4 mice per group. **(B)** Representative flow cytometry histograms showing the expression of activation markers CD40 and CD86 and checkpoint marker PD-L1 on pre-stimulated BMDCs, gating on live $CD11b^{+}CD11c^{+}$ cells.

### Figure S7

A

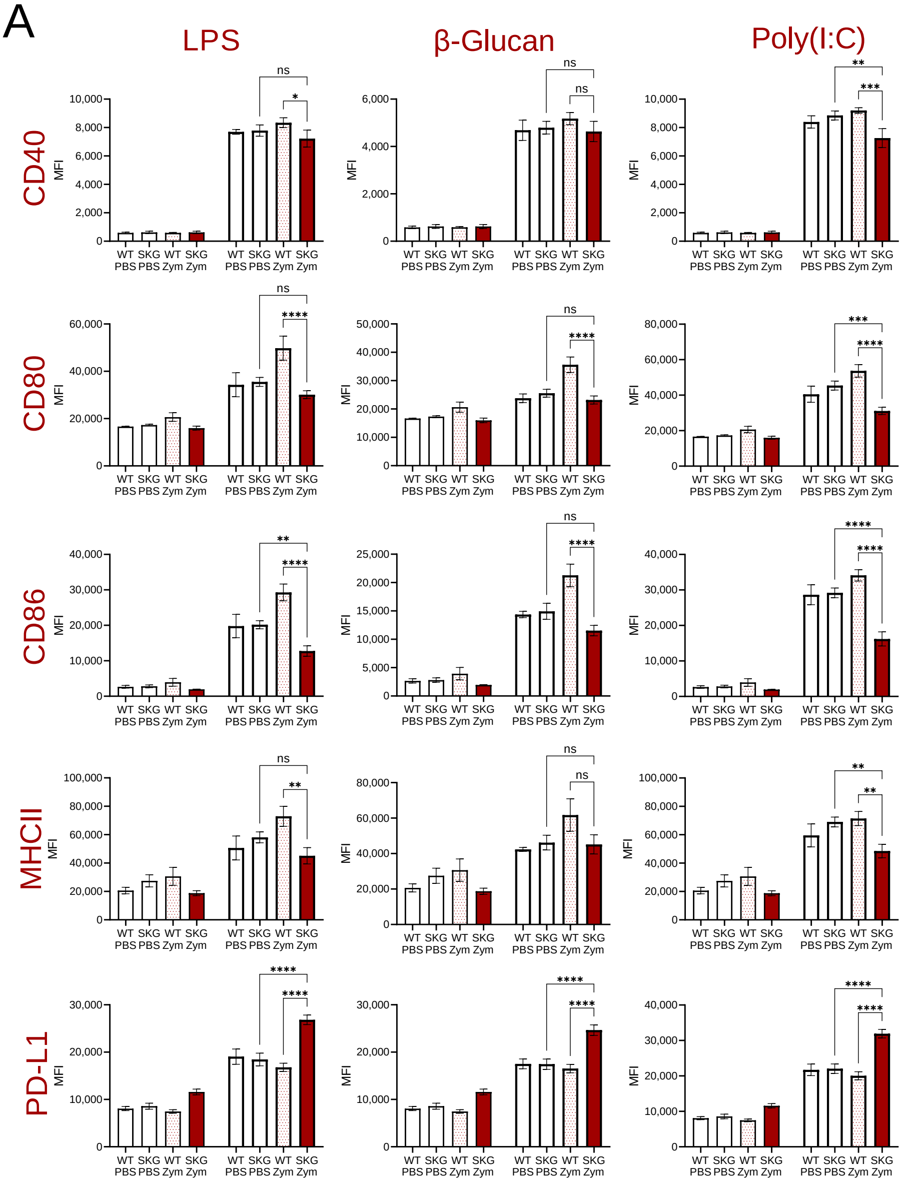

LPS
β-Glucan
Poly(I:C)
CD40
CD80
CD86
MHCII
PD-L1
MFI
WT SKG WT SKG WT SKG WT SKG
PBS PBS Zym Zym PBS PBS Zym Zym
ns
*
**
***
****


B

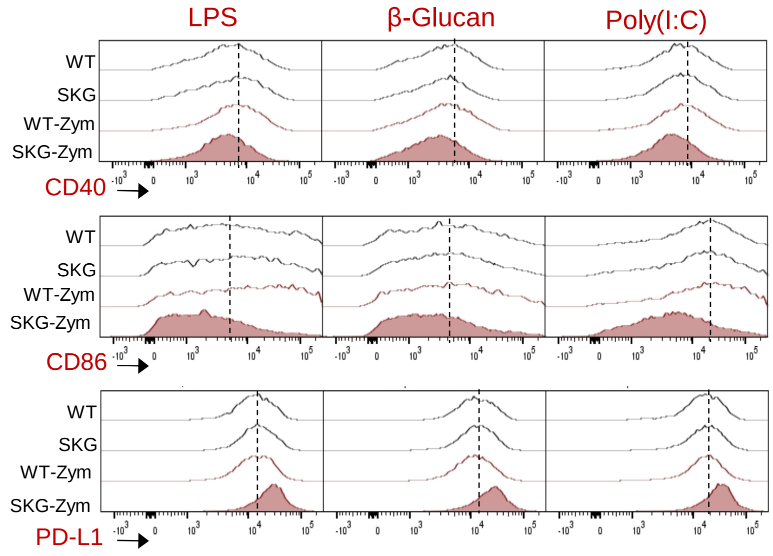

LPS
β-Glucan
Poly(I:C)
WT
SKG
WT-Zym
SKG-Zym
CD40
CD86
PD-L1

**Figure S8. Altered functional profile of dendritic cells derived *ex vivo* from the bone marrow of mice with active arthritis disease: collagen induced arthritis model (CIA).** BMDCs were derived *ex vivo* from the bone marrow of mice with collagen induced arthritis (CFA+Col RA) and control mice. BMDCs were stimulated with LPS 10 ng/mL, β-glucan 5 μg/mL, or poly(I:C) 20 μg/mL for 16 hours. **(A)** Expression of activation and checkpoint markers on BMDCs, analyzed by flow cytometry, pre-gating on live $CD11b^{+}CD11c^{+}$ cells; MFI – mean fluorescence intensity. Bars represent means ± SEM; statistical analyses with one-way ANOVA and Sidak's post-hoc test; * $p<0.05$, ** $p<0.01$, *** $p<0.001$; n=2-3 mice per group. **(B)** Representative flow cytometry histograms showing the expression of activation and checkpoint markers on pre-stimulated BMDCs.

### Figure S8

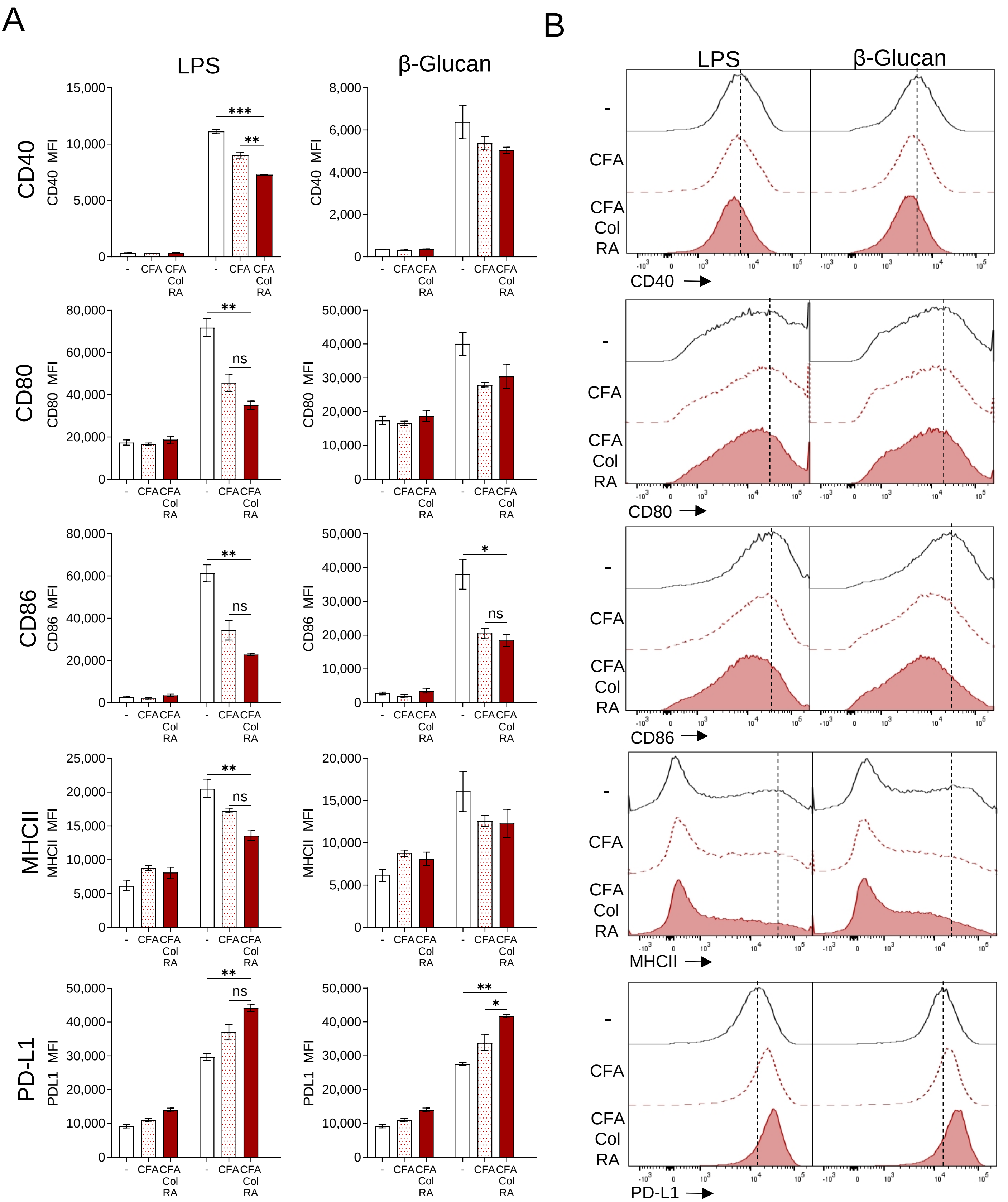
A
LPS
β-Glucan
CD40
CD40 MFI
CD80
CD80 MFI
CD86
CD86 MFI
MHCII
MHCII MFI
PD-L1
PDL1 MFI
-
CFA
CFA Col RA
***
**
*
ns
B
LPS
β-Glucan
-
CFA
CFA Col RA
CD40
CD80
CD86
MHCII
PD-L1

**Figure S9. Analyses of RA-SKG derived dendritic cells in chimeric mice. (A)** Relative abundance of $CD45.2^+$ RA-SKG versus control $CD45.1^+$ DCs in the chimeric mice, including cDCs in the mouse spleens and BMDCs derived in culture from the marrow of the chimeric mice. Relative percentages of cells in the $CD45.2^+$ versus $CD45.1^+$ gates are shown on the plots as means ±S.D. for all mice in each group. **(B)** Setting the gates on $CD45.2^+$ versus $CD45.1^+$ DCs using splenocytes and BMDCs from control CD45.2 and CD45.1 mice, stained, analyzed, and gated as the samples from the chimeric mice. **(C)** Expression of PD-L1 on BMDCs derived from the marrow of the chimeric mice, comparing $CD45.2^+$ RA-SKG and control $CD45.1^+$ DCs. Bars represent means ± SEM; statistical analyses with Student's *t*-test, with * $p<0.05$; MFI – mean fluorescence intensity. In all the experiments, cDCs in mouse spleens were gated as live $Lin^-F4/80^-CD64^-CD11c^+MHCII^+B220^-PDCA1^-$ cell (as shown in Figure S2B) and BMDCs in cultures as live $CD11b^+CD11c^+$ cells (as shown in Figure S5E).

Figure S9

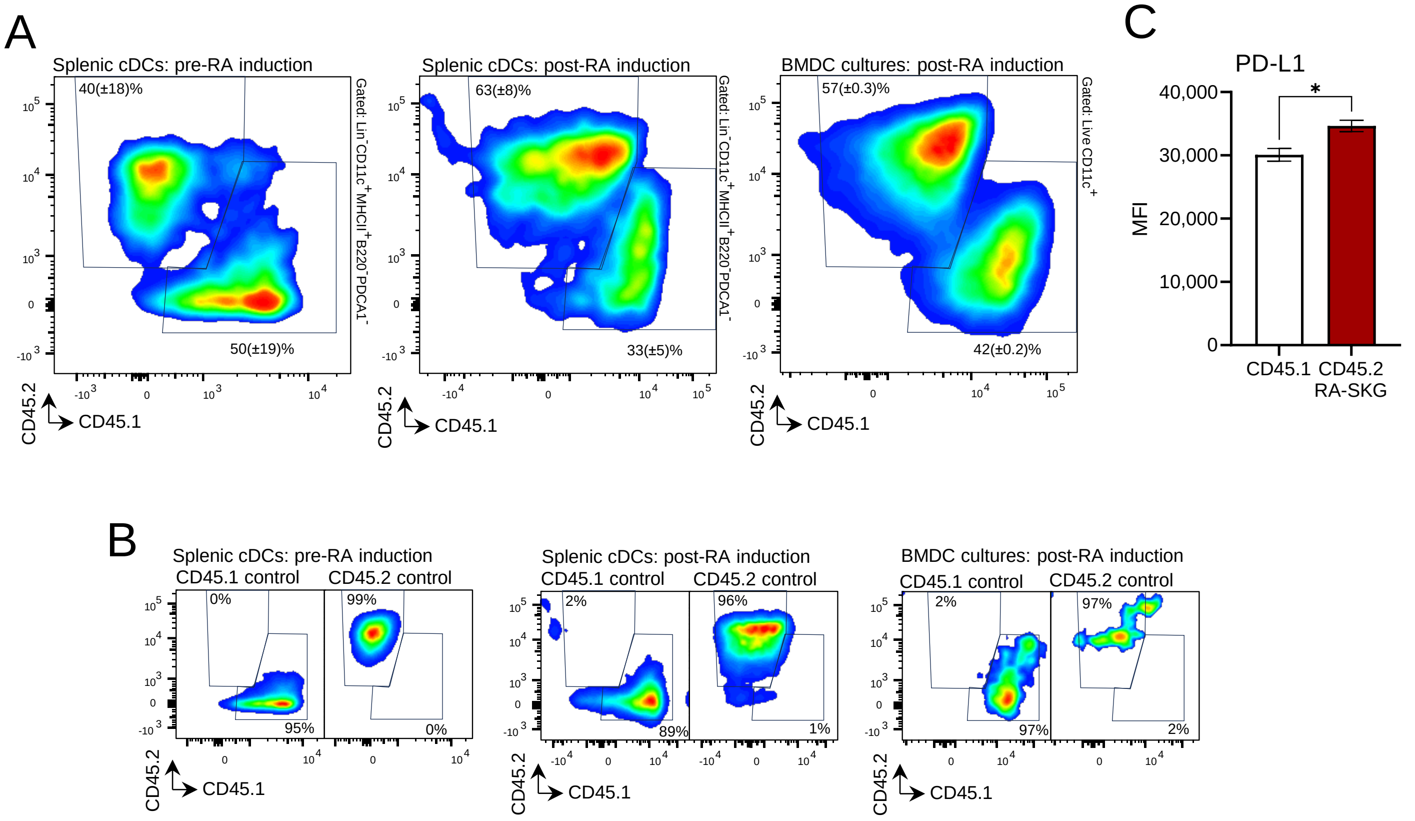

A
Splenic cDCs: pre-RA induction
40(±18)%
50(±19)%
Gated: Lin⁻CD11c⁺MHCII⁺B220⁻PDCA1⁻
CD45.2
CD45.1
Splenic cDCs: post-RA induction
63(±8)%
33(±5)%
Gated: Lin⁻CD11c⁺MHCII⁺B220⁻PDCA1⁻
CD45.2
CD45.1
BMDC cultures: post-RA induction
57(±0.3)%
42(±0.2)%
Gated: Live CD11c⁺
CD45.2
CD45.1
B
Splenic cDCs: pre-RA induction
CD45.1 control
CD45.2 control
0%
95%
99%
0%
CD45.2
CD45.1
Splenic cDCs: post-RA induction
CD45.1 control
CD45.2 control
2%
89%
96%
1%
CD45.2
CD45.1
BMDC cultures: post-RA induction
CD45.1 control
CD45.2 control
2%
97%
97%
2%
CD45.2
CD45.1
C
PD-L1
40,000
30,000
20,000
10,000
0
MFI
*
CD45.1
CD45.2
RA-SKG

**Figure S10. Altered gene expression profiles of DCs derived ex vivo from lineage depleted marrow of arthritis afflicted RA-SKG mice. (A-B)** Expression of genes encoding select cytokines, co-stimulatory molecules, and checkpoint markers in the RNA-seq data, presented as counts per million (CPM) and showing the genes with consistent changes in expression in RA-SKG versus control SKG DCs as compared to the previous analyses of the corresponding protein levels. Statistical analyses with one-way ANOVA and Sidak's post-hoc test; ** $p$<0.01, *** $p$<0.001, **** $p$<0.0001**. (C)** Heatmap displaying the genes significantly dysregulated in RA-SKG relative to control DCs, at FC > $|2.0|$ and FDR <0.001 under at least one of the conditions, corresponding to unstimulated, β-glucan stimulated, and poly(I:C) stimulated DCs. Gene expression is presented relative to the average of unstimulated control DCs. Hierarchical clustering of the gene expression signatures was performed using Pearson correlation and average linkage.

### Figure S10

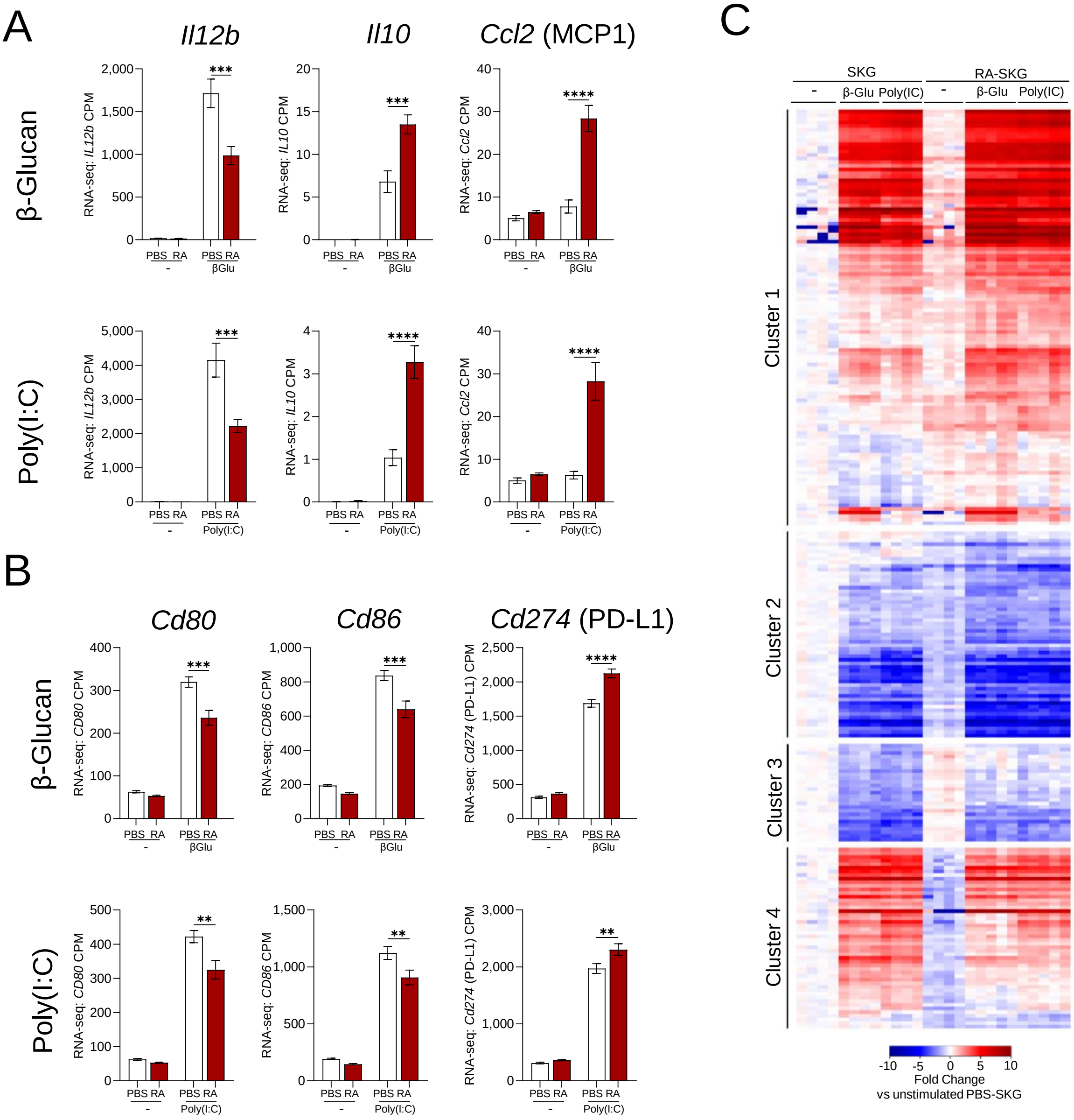
A
Il12b
Il10
Ccl2 (MCP1)
β-Glucan
Poly(I:C)
RNA-seq: IL12b CPM
RNA-seq: IL10 CPM
RNA-seq: Ccl2 CPM
PBS RA
-
βGlu
Poly(I:C)
***
****
B
Cd80
Cd86
Cd274 (PD-L1)
RNA-seq: CD80 CPM
RNA-seq: CD86 CPM
RNA-seq: Cd274 (PD-L1) CPM
**
C
SKG
RA-SKG
β-Glu
Poly(IC)
Cluster 1
Cluster 2
Cluster 3
Cluster 4
-10 -5 0 5 10
Fold Change
vs unstimulated PBS-SKG

**Figure S11. Monocyte derived DCs from rheumatoid arthritis patients: analyses of activation and responses to microbial stimulation. (A)** Confirmation of the successful derivation of moDCs over 8 days of culture of monocytes with GM-CSF and IL-4, based on CD11c and DC-SIGN marker expression. **(B-C)** Analyses of PD-L1 expression on moDCs of arthritis patients (RA) and matched healthy controls (HC), either at steady-state or following 16-hour stimulation with LPS 10 ng/mL, β-glucan 5 μg/mL, or poly(I:C) 20 μg/mL. MFI – mean fluorescence intensity. **(D)** Levels of IL-12 p40 in the culture supernatants of moDCs derived from arthritis patients (RA) and matched healthy controls (HC), at 16 hours of stimulation with β-glucan 5 μg/mL. **(E)** Levels of select cytokines and chemokines in the culture supernatants of unstimulated moDCs derived from arthritis patients (RA) and matched healthy controls (HC). Statistical analyses using paired *t*-test, * $p<0.05$; RA – samples from patients with rheumatoid arthritis; HC – samples from healthy controls, matched by age (±10 years) and sex.

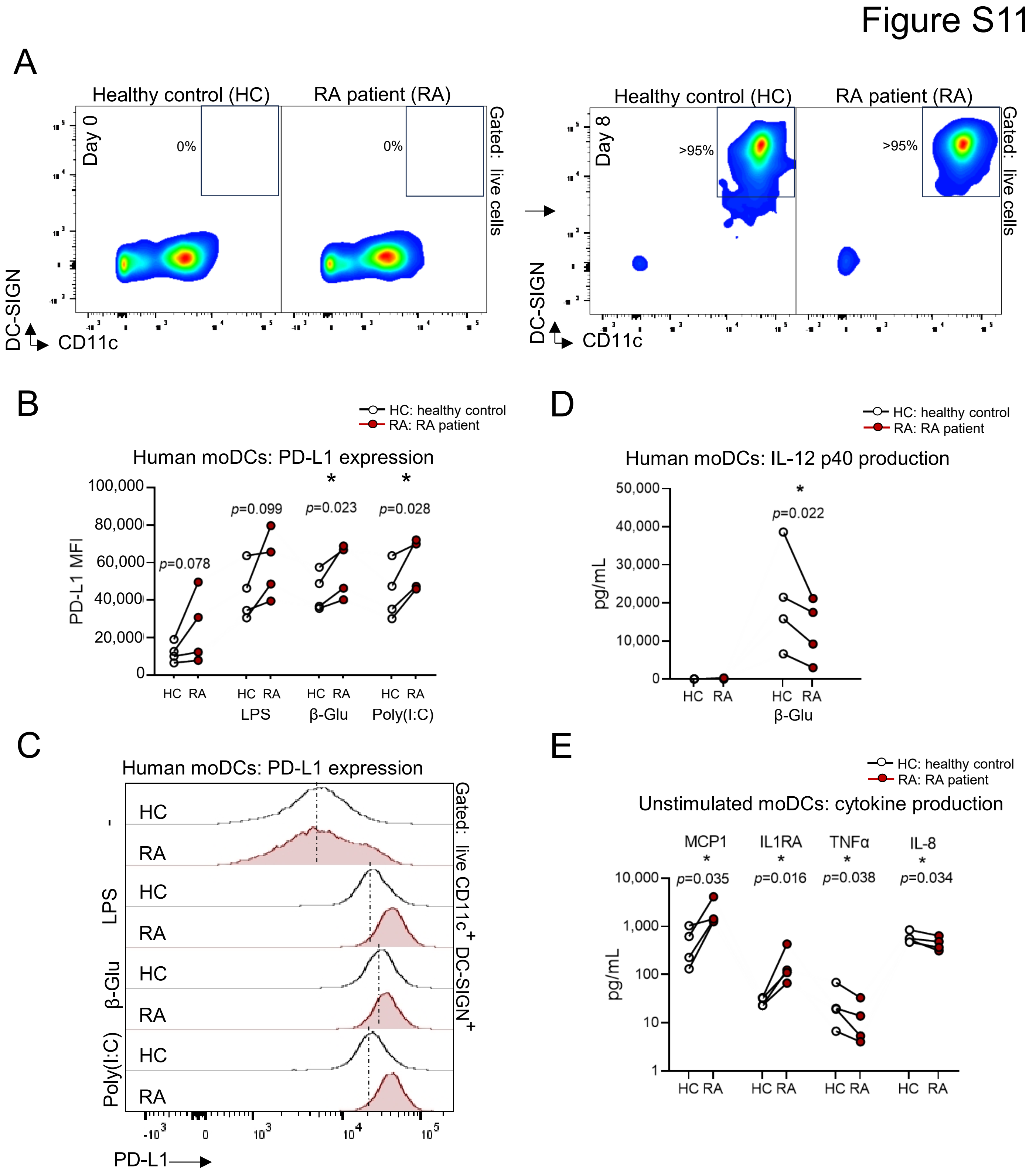

Figure S11
A
Healthy control (HC)
RA patient (RA)
Day 0
0%
0%
Gated: live cells
Day 8
>95%
>95%
DC-SIGN
CD11c
B
HC: healthy control
RA: RA patient
Human moDCs: PD-L1 expression
PD-L1 MFI
p=0.078
p=0.099
p=0.023
p=0.028
*
*
HC RA
LPS
β-Glu
Poly(I:C)
C
Human moDCs: PD-L1 expression
HC
RA
-
LPS
β-Glu
Poly(I:C)
Gated: live CD11c+ DC-SIGN+
PD-L1
D
Human moDCs: IL-12 p40 production
pg/mL
p=0.022
*
E
Unstimulated moDCs: cytokine production
MCP1
IL1RA
TNFα
IL-8
p=0.035
p=0.016
p=0.038
p=0.034

**Figure S12. Cell intrinsic impact of arthritis disease milieu on the capacity of murine DCs for T cell activation.** Analyses of OT-II CD4 T cell activation in co-cultures with DCs derived ex vivo from arthritis afflicted SKG-RA or control SKG-PBS mice, pre-stimulated with cognate antigen ovalbumin (OVA), together with LPS 10 ng/mL, β-glucan 5 μg/mL, or poly(I:C) 20 μg/mL for 16 hours. **(A)** Expression of cytokines IFNγ, IL-4, IL-17A, and IL-10 in live $CD3^{+}CD4^{+}$ T cells in the co-cultures with SKG-RA versus control SKG-PBS DCs, analyzed by flow cytometry and shown as the MFI of cytokine staining. Bars represent means ± SEM; statistical analyses with one-way ANOVA and Sidak's post-hoc test; * $p<0.05$, ** $p<0.01$, *** $p<0.001$; n=5-12 mice per group, consolidated from two independent experiments. **(B)** Representative flow cytometry plots showing activated $CD44^{hi}$ $CD62L^{lo}$ cells among live $CD3^{+}CD4^{+}$ T cells in the co-cultures with SKG-RA and control SKG-PBS DCs. The average percentage of cells in the gate for all mice in each group is indicated. **(C)** Quantification of activated $CD44^{hi}$ $CD62L^{lo}$ cells as a percentage of live $CD3^{+}CD4^{+}$ T cells in the co-cultures with SKG-RA and control SKG-PBS DCs. **(D)** Expression of CD44 and CD69 T cell activation markers, CD62L marker of naive T cells, CTLA4 marker of T cell exhaustion, and Ki67 marker of cell proliferation, gating on live $CD3^{+}CD4^{+}$ T cells in the co-cultures with SKG-RA and control SKG-PBS DCs. MFI – mean fluorescence intensity. **(E)** Representative flow cytometry histograms showing the expression of CD44, CD62L, CD69, CTLA4 and Ki67 on/in live $CD3^{+}CD4^{+}$ T cells in the co-cultures with SKG-RA and control SKG-PBS DCs. **(B-E)** Bars represent means ± SEM; statistical analyses with one-way ANOVA and Sidak's post-hoc test; * $p<0.05$, ** $p<0.01$, *** $p<0.001$; n=5-6 mice per group.

### Figure S12

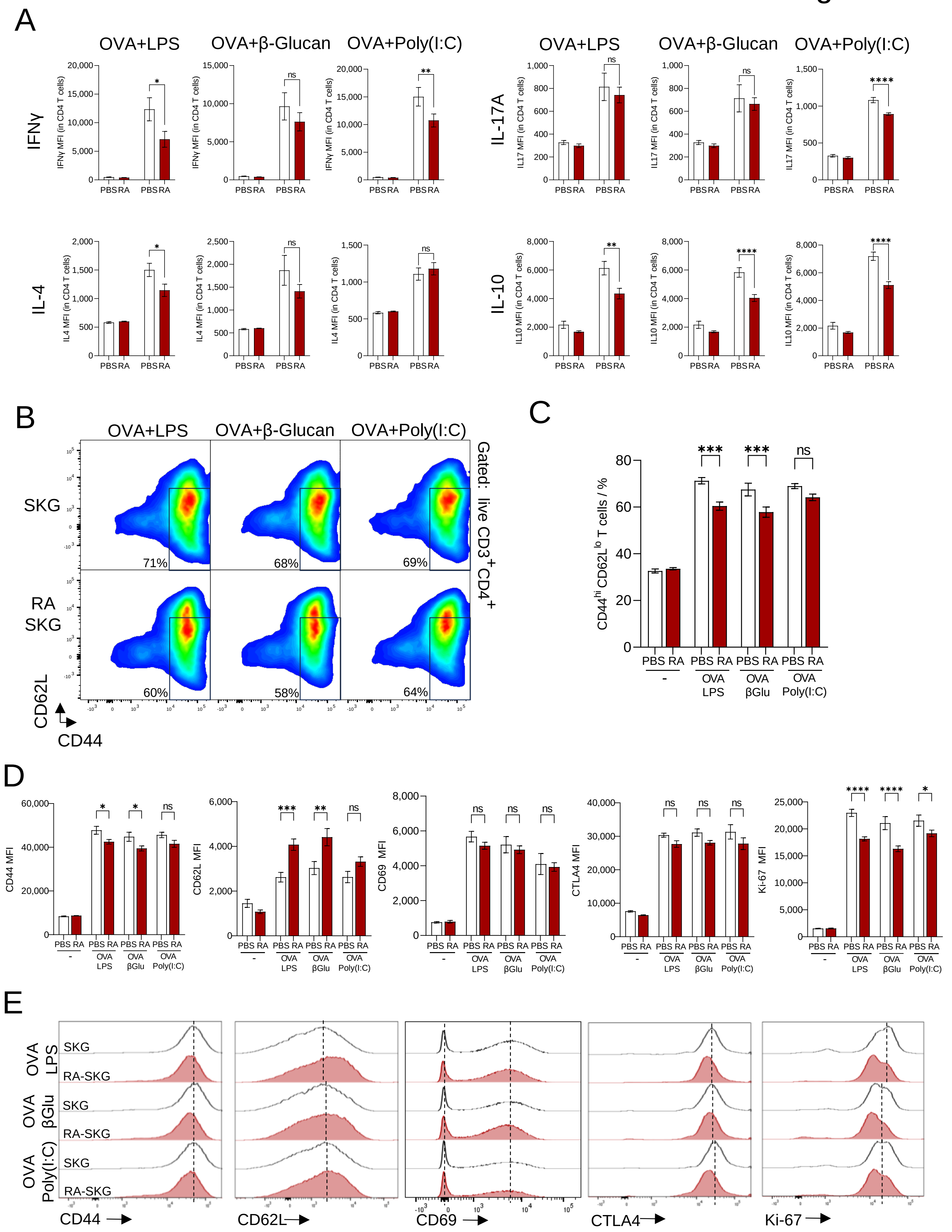

A
OVA+LPS
OVA+β-Glucan
OVA+Poly(I:C)
IFNγ
IL-17A
IL-4
IL-10
PBS RA
B
SKG
RA SKG
71%
68%
69%
60%
58%
64%
Gated: live CD3+CD4+
CD62L
CD44
C
CD44hi CD62Llo T cells / %
***
***
ns
OVA LPS
OVA βGlu
OVA Poly(I:C)
D
CD44 MFI
CD62L MFI
CD69 MFI
CTLA4 MFI
Ki-67 MFI
E
OVA LPS
OVA βGlu
OVA Poly(I:C)
SKG
RA-SKG
CD44
CD62L
CD69
CTLA4
Ki-67
